# The formation and closure of macropinocytic cups in a model system

**DOI:** 10.1101/2022.10.07.511330

**Authors:** Judith E. Lutton, Helena L. E. Coker, Peggy Paschke, Christopher J. Munn, Jason S. King, Till Bretschneider, Robert R. Kay

## Abstract

Macropinocytosis is a conserved endocytic process where cells take up medium into micron-sized vesicles. In *Dictyostelium*, macropinocytic cups form around domains of PIP3 in the plasma membrane and extend by actin polymerization. Using lattice light-sheet microscopy, we describe how cups originate, are supported by an F-actin scaffold and shaped by a ring of actin polymerization, created around PIP3 domains. How cups close is unknown. We find two ways: lip closure, where actin polymerization at the lip is re-directed inwards; and basal closure, where it stretches the cup, eventually causing membrane delamination and vesicle sealing. Cups grow as expanding waves of actin polymerization that travel across the cell surface, capturing new membrane. We propose that cups close when these waves stall. This ‘stalled wave’ hypothesis is tested through a conceptual model, where the interplay of forces from actin polymerization and membrane tension recreates many of our observations.

## Introduction

Cells must continuously transport material across their plasma membrane to sustain their life. Macropinocytosis is a conserved endocytic process in which cells project cups from their plasma membrane to engulf droplets of medium into micron-sized vesicles ^1-4^. It is used for feeding by cancer cells and amoebae ^5, 6^, and to take up antigens by immune cells ^7^. Viruses and bacteria use macropinocytosis to infect cells ^8^, while it is the delivery route for mRNA vaccines ^9^. Despite this clear medical importance, and its discovery nearly 100 years ago ^10^, only recently has macropinocytosis started to attract major attention ^11^. Fundamental questions therefore remain, such as: how are macropinocytic cups formed; what supports them; what makes them close; and how do they close?

Macropinocytosis is driven by the actin cytoskeleton, with PIP3 also playing a crucial role, as shown by genetic and inhibitor studies and its localization to forming macropinosomes ^12-15^. In *Dictyostelium*, PIP3 forms discrete domains in the plasma membrane, which can be microns across, and around which macropinocytic cups form ^16, 17^. These domains are complex and include active-Ras, which activates PI3-kinase, and active-Rac, which activates actin polymerization ^18^. Domains are regulated by RasGAPs such as NF1 and RGBARG ^19-21^, while PIP3 itself recruits effectors required for macropinocytosis, such as the protein kinase Akt and actin polymerization regulator, Leep1 ^22, 23^. PIP3 domains form cell-autonomously, without the need for external signals. How they function in macropinocytosis remains unclear.

We have proposed that PIP3 domains recruit the Scar/WAVE complex to their periphery, thus creating a ring of protrusive F-actin around themselves and forming the walls of macropinocytic cups ^18^. While this property of PIP3 domains helps explain how cups are shaped, it does not explain how they close.

Although macropinocytic cups are readily visible by microscopy, they are hard to follow in 3D due to their size, dynamism and the light-sensitivity of cells. Lattice light-sheet microscopy (LLSM) gives a new approach, allowing macropinocytic cups to be followed in 3D from birth to closure ^18, 24-26^. To match this advance in microscopy, image analysis methods need extending to 3-dimensional time series, to segment cells and PIP3 domains and to measure fluorescent intensities along the plasma membrane ^27^.

Here, using lattice light sheet microscopy and custom computational analysis, we have followed macropinocytosis and PIP3 domains in the *Dictyostelium* model with unprecedented detail. We describe how cups are structured, expand and close, and infer some general principles behind their organization. This leads to a physical model, based on the behaviour of PIP3 domains, which reproduces cup closure and several other features that we observe.

## Results

### Macropinosomes form from cups that can close at lip or base

Lattice light-sheet microscopy allows macropinocytosis to be followed for 5-10 minutes in *Dictyostelium* cells in 3D at around 0.3 Hz, with two fluorescent channels. Cells were observed at low density in a simple medium, where they maintain macropinocytosis for at least a day ^28^. We initially used Ax2 cells with paired reporters for PIP3 (PkgE-PH-GFP), marking the membrane of macropinocytic cups, and F-actin (lifeAct-mCherry) revealing the actin cytoskeleton (**Figure 1; movie 1**).

**Figure 1:**
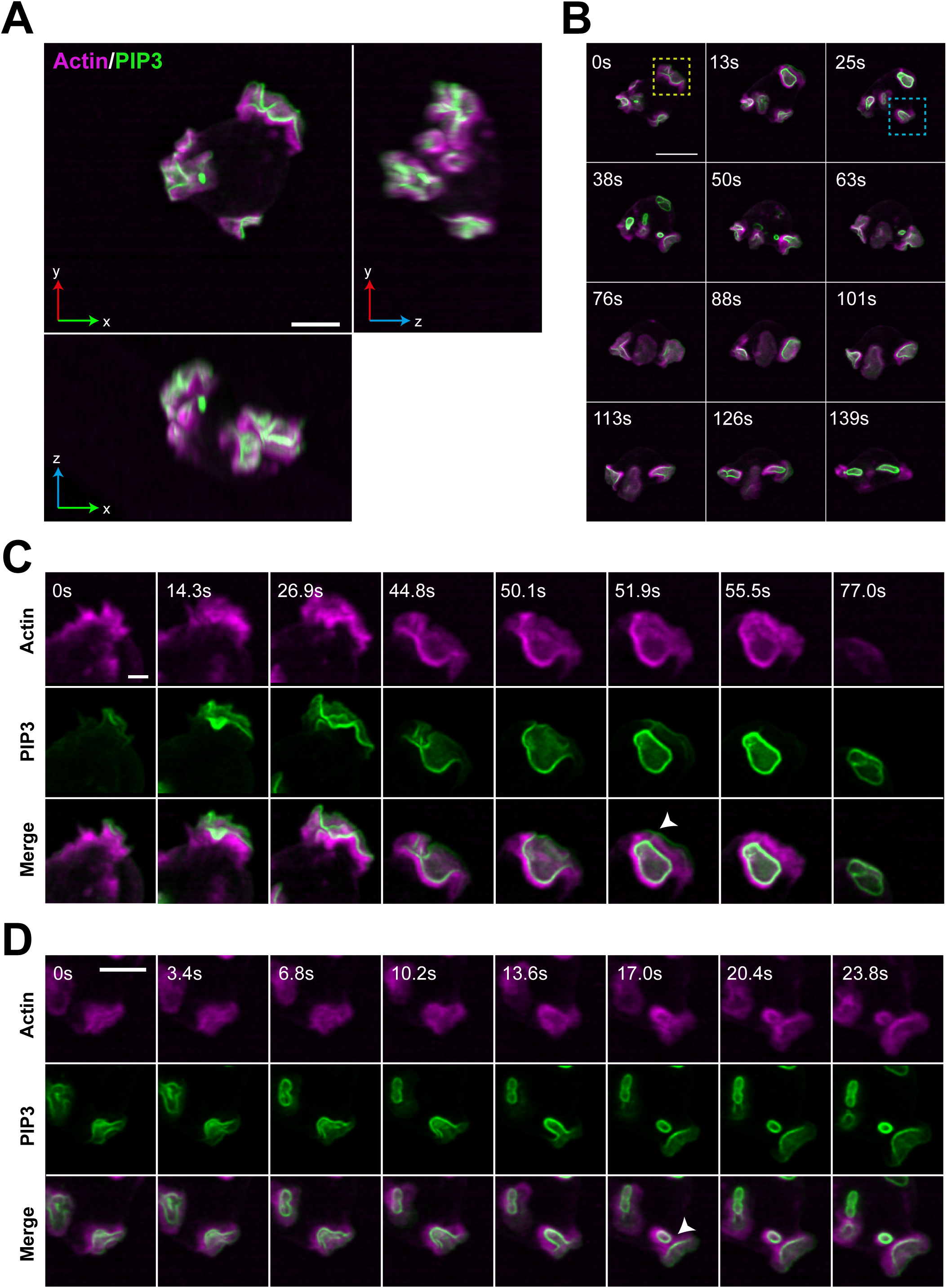
Lattice light-sheet microscopy shows that macropinocytic cups have two ways of closing. Images from a lattice light-sheet movie of a *Dictyostelium* cell expressing reporters for PIP3 (green) and F-actin (magenta). The PIP3 reporter reveals the PIP3 domain of macropinocytic cups and early macropinosomes and the F-actin reporter reveals their structural scaffold. (A) Orthogonal projections of a single cell (x,y top left) having three macropinocytic areas. (B) Time sequence from the same cell. Yellow-boxed area: cup closing at the lip releases a large macropinosome at 13 sec, whose PIP3 signal intensifies and then fades away by 50 sec. The domain itself is extinguished. Blue-boxed area: successive basal closures at a second PIP3 domain. Small macropinosomes are released at 38 sec and 63 sec, before the domain closes at the lip at 139 sec and is extinguished. (C) Lip-closure at higher temporal resolution with split colours, corresponding to the yellow-boxed area. Arrow: a tether between macropinosome and the cell surface revealed by the PIP3 reporter – it is presumed to be a thin tube. (D) Basal-closure at higher temporal resolution with split colours, corresponding to the blue-boxed area. Arrow: newly separated macropinosome and persistent PIP3 in plasma membrane. Ax2 cells transformed with pPI304 to express reporters for PIP3 (PkgE-PH-GFP) and F-actin (lifeAct-mCherry) were viewed in imaging medium by LLSM with full volumes taken every 1.7 sec. Maximum intensity projections are shown. Scale bars: A and B = 5 μm; C and D = 2 μm. See movies 1-3.

Macropinosomes in *Dictyostelium* cells form almost exclusively from cups projected from the cell surface, consistent with earlier work showing ‘crowns’ on growing cells ^29 6^. *Dictyostelium* cells thus resemble macrophages ^15^, though we did not observe macropinosomes forming from linear ruffles ^26^, nor were ‘tent-pole’ filopodial projections of F-actin ^25^ notably involved.

Cups form on any exposed part of cells and frequently move around as they develop. On average cells have 2.71±1.71 PIP3 domains and produce 1.8±1.2 macropinosomes per minute **(Table S1)**, ranging up to more than 5 microns in their longest dimension. Rates and sizes varied greatly. After closing, macropinosomes lose first their F-actin coat, and then their PIP3 reporter.

#### Macropinocytic cups can close in two different ways

Unexpectedly, we observed that macropinocytic cups can close in two ways, which we call lip and basal closures, of which lip closure is roughly twice as common (**Figure 1, Table S1, movies 1-3**). In lip closure, cups close at, or near, the lip and include nearly all the PIP3 domain in the resulting macropinosome. Residual PIP3 in the plasma membrane fades away, extinguishing the domain and terminating macropinocytosis from that site. In a complex variant, macropinocytic cups form lobes, which appear as over-lapping arcs of PIP3-reporter in maximum-intensity projections. When these composite structures close, several macropinosomes are released almost simultaneously (**Figure S1A and B**; **movie 4**).

In basal closure, cups deepen and narrow, then constrict locally before pinching off a macropinosome near the base. A large proportion of the PIP3 domain remains in the plasma membrane, where it can yield further macropinosomes. This sequence generally terminates by closure at the lip and extinction of the PIP3 domain.

Macropinosomes formed by either route frequently remained attached to the cell surface by a thin tether, which can persist for 10 seconds or more, before rupturing (**Figure 1C**, 51.9 sec; also **Figure S1A** and **B**. As tethers are visible with the PIP3 reporter, we assume that they are narrow tubes of membrane, although the individual walls are not resolved. This implies that macropinosomes are not fully sealed and released until the tether is ruptured. Tethers are under further investigation.

Cups can also fail. Most commonly, the PIP3 domain simply fades away, followed by its associated F-actin. Or there can be an abortive closure in which the cup constricts but does not close (**Figure S1C, Movie 5**).

#### Macropinocytosis is similar in non-axenic cells

Strain Ax2 is a laboratory workhorse, which like other axenic strains has been selected for growth in liquid medium. This is enabled by a high rate of macropinocytosis due to deletion of the conserved RasGAP, NF1 ^19^. We therefore asked whether non-axenic cells, with intact NF1, perform macropinocytosis in a similar way.

DdB cells ^30^ were grown in enriched liquid medium to stimulate macropinocytosis and examined using the PIP3/F-actin reporter combination. They produce generally smaller PIP3 domains and macropinosomes than Ax2 cells (**Figure S1D-G)**. Both lip and basal closures, as well as tethers, are observed, showing that these are common features of *Dictyostelium* macropinocytosis (**Movies 6 and 7**).

### How macropinocytic cups are organized: PIP3 domain and F-actin scaffold

PIP3 domains are singular features of macropinocytic cups in *Dictyostelium* ^14, 17^. To use these as landmarks for mapping other components, we developed computational tools to identify PIP3 domains by first segmenting the cell surface and then the PIP3 domains themselves (**Figure 2A**). This two-step procedure means that PIP3-rich vesicles are excluded, once released from the cell surface. Intensities of a panel of reporters and GFP-fusions (see methods and **Table S3)** were then measured along the membrane from the domain boundary, with the bottom of the cup providing a second reference point, and geometric parameters calculated, such as domain area and curvature. Importantly, this approach allowed us to average over multiple events.

**Figure 2:**
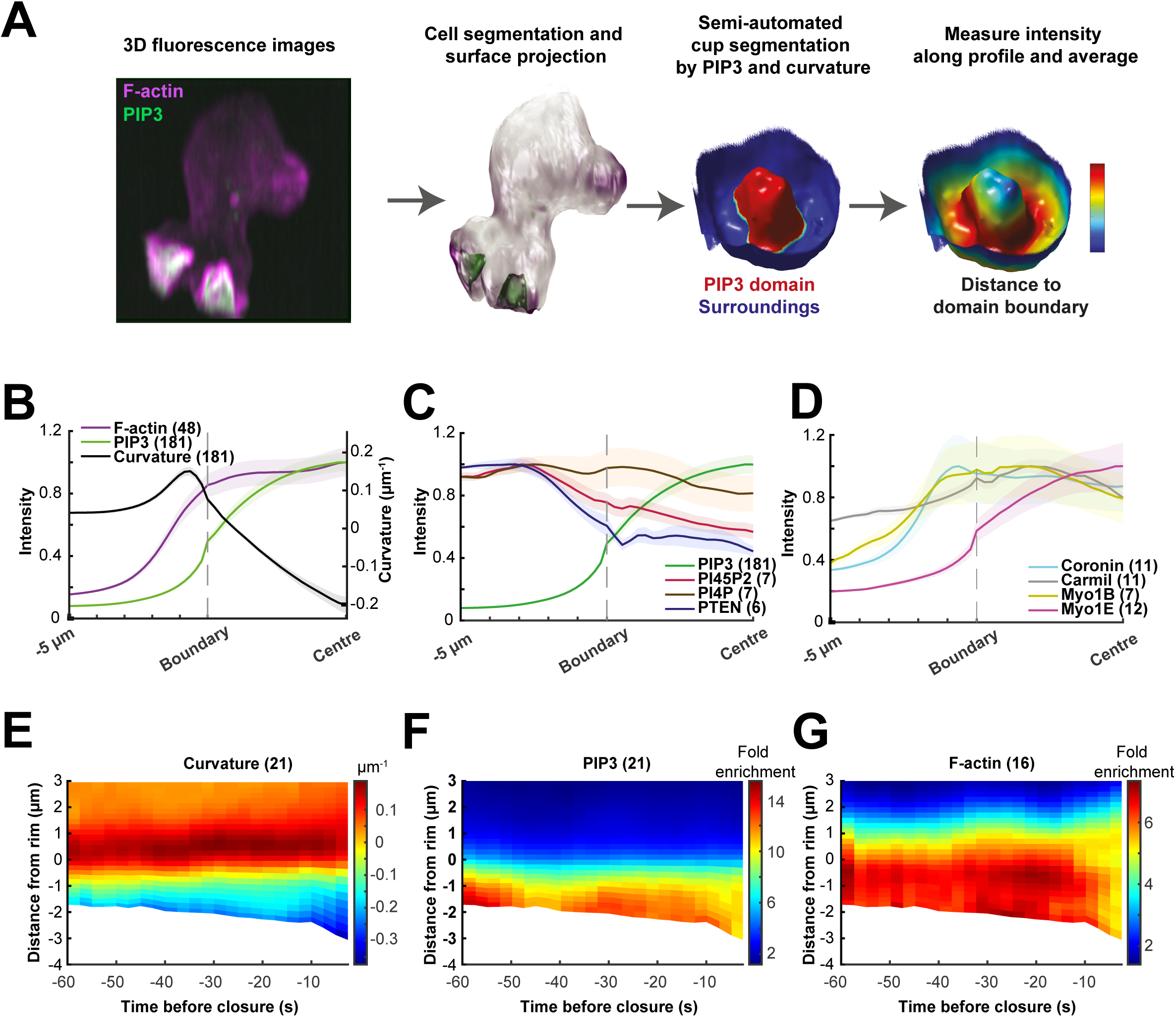
Macropinocytic cups are organized around PIP3 domains and supported by F-actin scaffolds. (A) Mapping strategy: PIP3 domains were defined in a two-step process, yielding first the cell surface segmented by a 3D method using both fluorescent reporters; and then the PIP3 domains, segmented using the PIP3 reporter, and including a curvature correction for spill-over across membrane folds. Domains were inspected and annotated manually using virtual reality. Vesicles not attached to the plasma membrane are excluded by this process. The lip of cups is defined by the inflection in membrane curvature. (B) PIP3 domain and F-actin scaffold: the PIP3 domain occupies the inner surface of the cup, with little spill-over beyond the lip, and the cup is supported by a continuous F-actin scaffold from bottom to just beyond the lip. (C) Cytoskeletal proteins: coronin, carmil (modestly), and myosin1B and 1E are all concentrated in macropinocytic cups, with the PIP3-binding myosin1E mirroring PIP3 and coronin concentrated just beyond the lip. (D) Phosphoinositide gradients: PIP3 is strongly graded with its peak towards the centre of the domain, whereas PTEN, which breaks PIP3 down, is excluded from PIP3 domains. PI4,5P2, the substrate for PIP3 production, shows an outside-in gradient. This is consistent with PIP3 being produced within PIP3 domains and diffusing out for destruction by PTEN; whereas PI4,5P is depleted within the domain by PIP3 formation and replenished by diffusion from outside. PI4P shows little variation. In (B,C,D) the domain boundary is set as zero and distances into the domain are negative and normalized, whereas those outside it are actual distances. (E, F, G) Space-time plots over the last minute before macropinocytic cups close. Both lip and basal closures are included. Cups deepen particularly in the last 10 sec before closure, as shown by the increasingly negative distance from the boundary. There is evidence for increased accumulation of PIP3 and F-actin at 10 to 40 sec before closure. In (E,F,G), the PIP3 domain boundary is set as zero with negative numbers being into the domain and positive beyond it. In all cases the bracketed numbers are the number of cups analysed.

#### PIP3 domains occupy the inner face of macropinocytic cups and produce phosphoinositide gradients

We define the lip of cups by the inflection in their membrane curvature and find that the PIP3 domain boundary falls just inside the lip, with PIP3 levels elevated over the entire inner face of the cup and falling off further out (**Figure 2B**). Mapping the distribution of PIP3 shows a striking gradient within domains, with its high point at their base.

PI4,5P2 - the substrate for synthesis of PIP3 by PI3-kinases – has a reciprocal gradient with levels lower inside macropinocytic cups than outside (**Figure 2C**). PTEN, which reverts PIP3 to PI4,5P2, also has a reciprocal distribution, with high levels outside the cup and over the whole plasma membrane, and low levels within the cup, consistent with previous reports ^31^.

PI4P, a relatively abundant phosphoinositide of the plasma membrane, shows little change in distribution in PIP3 domains, while PI3,4P2 is near background, but increases strongly in macropinosomes once cups close (not shown).

#### PIP3 domains are critical for macropinocytosis

To help define the importance of PIP3 domains in macropinocytosis, we examined mutants of PIP3 signalling (**Figure S2)**. These have greatly reduced fluid uptake ^14, 18^, but their functional defects have been hard to pin down.

At one extreme, in a mutant lacking all five ‘type-1’ PI3-kinases ^32^, PIP3 domains are abolished, but the underlying domains of active Ras still form. These appear incapable of macropinocytosis. Conversely, when PTEN is deleted the PIP3 domains grow to encompass the entire cell surface but are capable of only very inefficient macropinocytosis, forming occasional small macropinosomes. A similar, but less extreme phenotype is produced by deletion of the RasGAP, RGBARG in which cells are only able to perform base closures^20^. Thus, PIP3 domains are essential, or nearly so, for macropinocytosis, but must be properly regulated.

#### Macropinocytic cups are mechanically supported by an F-actin scaffold

Macropinocytic cups require structural support to prevent their collapse by membrane tension. We therefore examined their actin cytoskeleton using reporters for selected cytoskeletal proteins, each paired with a PIP3 reporter (**Figure 2D, Figure S3**). No clear structural differences were observed between cups that ultimately closed at lip or base, allowing inclusion of both in our analysis.

This mapping shows that macropinocytic cups have a continuous layer of F-actin over their whole extent, from top to bottom, which is very clear in deep cups and extends some distance beyond the rim (**Figure 2B; see also Figure 1B**,**C**). This layer is thicker in the cup than in more remote areas of the plasma membrane, consistent with it providing structural support.

Coronin binds F-actin, with a preference for newly formed filaments and regulates actin dynamics ^29 33^. It is strongly recruited to cups, concentrating towards the rim. Single-headed myosins link F-actin to membranes ^34^, with several strongly recruited to macropinocytic cups ^35^. Myosin-1B is recruited towards the periphery of cups, whereas the PIP3-binding myosin-1E is concentrated towards the base, closely matching the distribution of PIP3. Carmil ^36^ is only modestly enriched in cups. Although microtubules (labelled with GFP-alpha-tubulin) moved actively within the cytoplasm and made fleeting contacts with macropinocytic structures, we did not detect any prolonged association between them suggestive of a structural role in cups.

These results show that macropinocytic cups are organized around essential PIP3 domains and supported by a specialized F-actin scaffold, in which the myosin-1 proteins may be linkers to the membrane that prevent it delaminating.

### Evidence that PIP3 domains shape rings of dendritic actin polymerization around themselves

We have proposed that PIP3 domains produce cupped structures by creating a ring of actin polymerization around their periphery ^18^. To further test this idea, we took advantage of the sensitivity of LLSM to image the Scar/WAVE complex – which activates dendritic actin polymerization - through the entire life cycle of macropinocytic cups. **Figure 3** and **Movies 8 and 9** show that NAP-GFP is recruited in striking rings around PIP3 domains. A reporter for the Arp2/3 complex, which is activated by Scar/WAVE, was also recruited in a similar manner (**Figure 3B, Movie 10**), as also recently reported ^23^.

**Figure 3:**
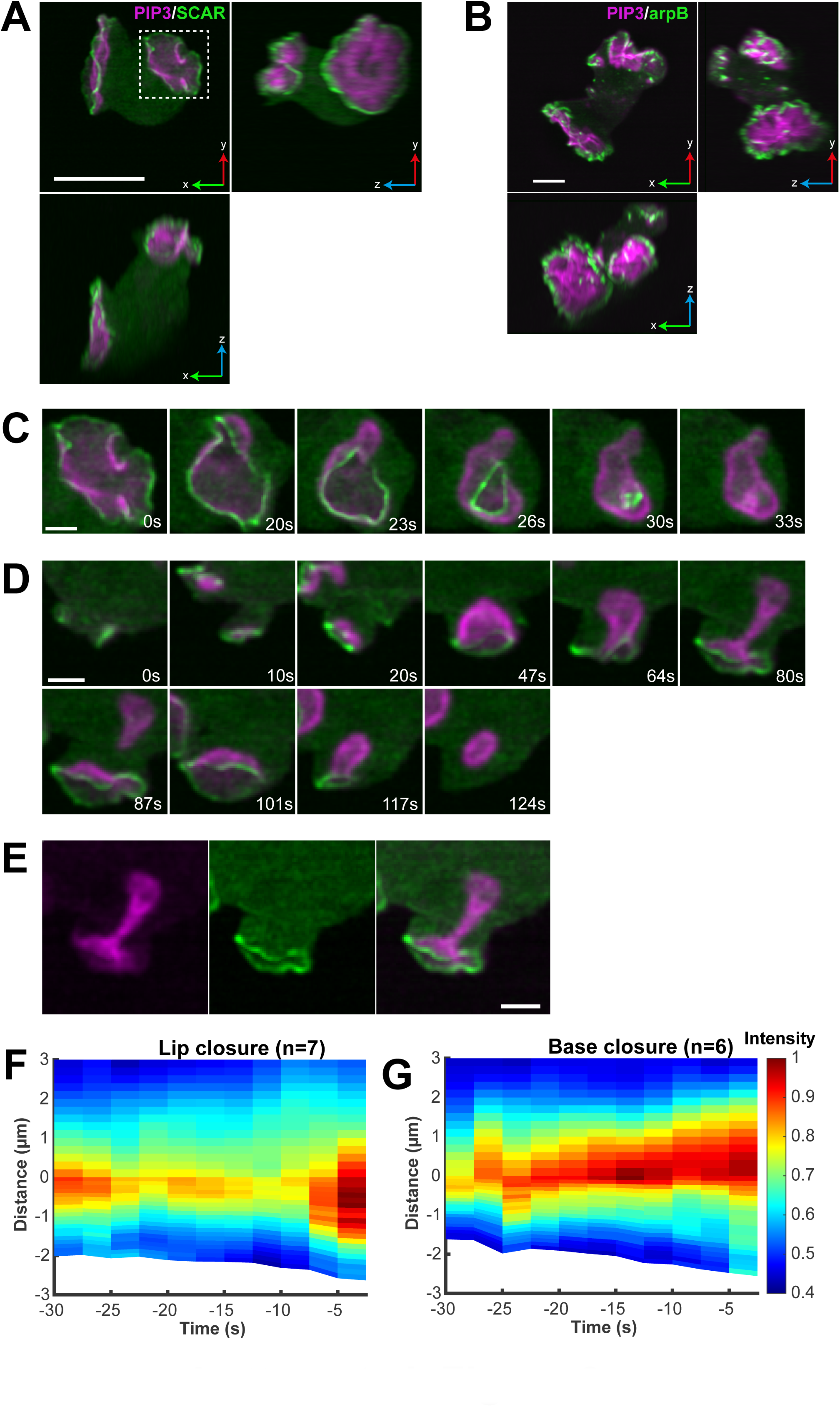
PIP3 domains shape cupped structures by attracting a ring of dendritic actin polymerization to their periphery. Dendritic actin polymerization is initiated by the Arp2/3 complex, which is activated by the Scar/WAVE complex. (A) Orthogonal projections of a single cell with two macropinocytic cups, expressing reporters for PIP3 (magenta) and the NAP subunit of the Scar/WAVE complex (green). (B) Orthogonal projections of a single cell with two macropinocytic cups expressing reporters for PIP3 (magenta) and the Arp2 subunit of the Arp2/3 complex (green). (C) En face view of a cup closing at the lip, showing a contracting ring of Scar/WAVE, indicating that closure is driven by actin polymerization. (D) A cup producing macropinosomes by basal closure (87 sec) and lip closure (124 sec). Scar/WAVE is present at the lip of the cup throughout, but not at the site of the basal closure. Lip closure extinguishes the domain. (E) Split channels of the 80 sec panel of (D). (F,G) Space/time plots showing the distribution of Scar/WAVE in cups closing either at the lip or base showing the continuous presence of Scar/WAVE at or near the lip of the cup. Closure occurs at 0 sec and the edge of the PIP3 domain is set at zero, with negative numbers representing distances inside the cup. Scale bars: A and B = 5 μm; C-D = 2 μm. See movies 8-10.

Following the NAP-GFP rings over time demonstrated that they are ever-present at the rim of cups, from early in their life, through their expansion and into closure. The persistence of Scar/WAVE at the edge of PIP3 domains was quantified and confirmed by space-time plots (**Figure 3F and G**). Taking recruitment of the Scar/WAVE and Arp2/3 complexes as reporting sites of actin polymerization ^18^, these results confirm that PIP3 domains are encircled by a ring of dendritic actin polymerization.

### How cups close

En face views show that lip closure occurs by concerted inward movement of the lips until the orifice is sealed (**Figure 3C, Movie 11**). In principle, this might be due to a purse-string constriction, driven by myosin-II, as proposed for phagocytic cups ^37^. However, we could not detect myosin-II at the lip or elsewhere in cups (**Figure S3**) and macropinocytosis is insensitive to blebbistatin inhibition ^28, 38^. Rather, the Scar/WAVE reporter is present as a contracting ring throughout closure. Thus, we propose that cups close by actin polymerization at their lip, which is re-directed inwards.

Examining basal closures in a similar way, we found that Scar/WAVE and Arp2/3 were not detectably recruited at the site of closure. This is shown in **Figure 3D** (**Movie 9**), where a basal closure is followed by a lip closure. Scar/WAVE is not recruited to the basal closure site, but only to the lip, providing a positive control for the reporter. This indicates that base closure is not driven by localized actin polymerization at the site of closure.

Significantly, Scar/WAVE remains at the lip of cups during basal closure, suggesting that actin polymerization continues there, and may drive closure indirectly. Accordingly, the cup elongates and narrows (**see Figure S4**), presumably stretching the membrane, until eventually a macropinosome is pinched off.

Consistent with this, mapping F-actin and PIP3 through the last minute of cup lifetime shows that both reporters maintain their relative distributions, possibly with some intensification 20-30 seconds before closure (**Figure 2E-G**).

Detailed observation of basal closure shows that the PIP3-positive membrane often delaminates from the F-actin scaffold at the site of closure (**Figure 4 Movies 11-13**). This indicates both that the membrane is under significant tension and that local pinching by the actin cytoskeleton is not the cause of closure. Similar delaminations are also observed with cups closing near the lip, indicating that membrane tension also contributes to lip closure.

**Figure 4:**
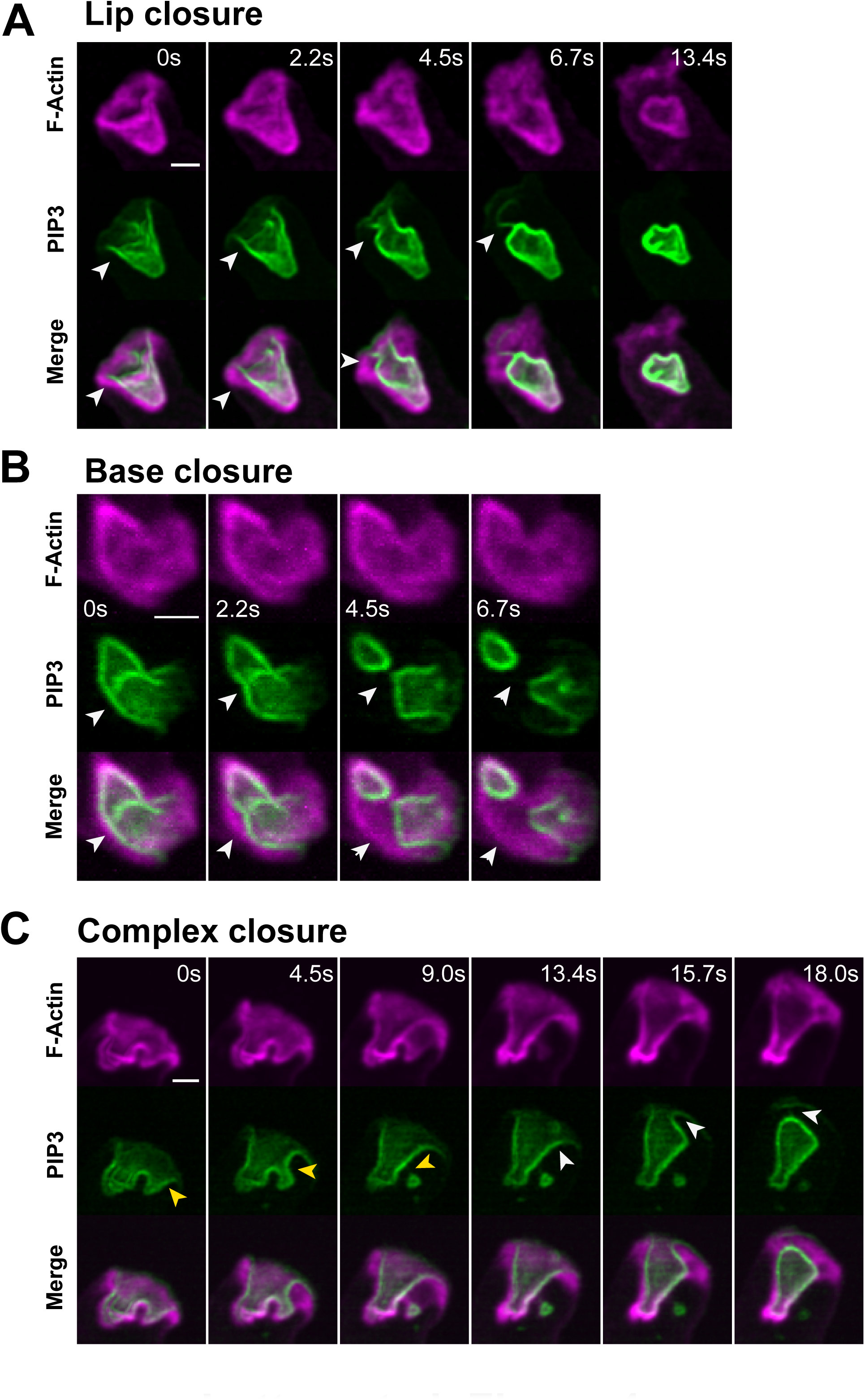
Delamination during cup closure: evidence that the membrane within cups is under tension. By delamination we mean the separation of the plasma membrane from the underlying actin cytoskeleton and take this as evidence that the membrane is under tension. Delamination most usually occurs as a macropinosome is sealing. (A) Closure just below the lip with delamination arrowed. Note also the tether at 6.7 sec. (B) Base closure with delamination arrowed. (C) Complex closure with delamination arrowed. Ax2 cells expressing reporters for PIP3 (PkgE-PH-GFP) and F-actin (lifeAct-mCherry). Scale bars = 2 μm. See movies 11-13.

Superficially, the two closure methods appear to have different mechanisms, but both involve continued actin polymerization at the lip, driven by Scar/WAVE. We propose that cups close at the lip by redirecting this actin polymerization inwards; and at the base by using it to deepen the cup, increasing membrane tension until delamination and basal sealing of a macropinosome occurs.

### Macropinocytic cups close when their PIP3 domain ceases expanding

What triggers macropinocytic cup closure? We noticed from area measurements that cups tend to close when their PIP3 domain stops expanding. Since this leads to a plausible model for closure (next section), we examined the correlation in detail.

First, we analysed a set of domains that could be followed from their origin to the formation of a macropinosome by lip closure (**Figure 5A and B, Movie 14**). All originate with a small PIP3 domain associated with a pre-existing focus of F-actin (11/11 cases). This F-actin focus is usually small, but occasionally can be a pseudopod, which is then subverted into a macropinocytic cup. The individual traces of domain area, perimeter and depth are strikingly variable, whether aligned by domain initiation or closure (**Figure 5C and D**) due primarily to their spread of lifetimes (30-130 sec). To better discern overall trends, we combined plots by normalizing both time and the geometric parameters (**Figure 5E**).

**Figure 5:**
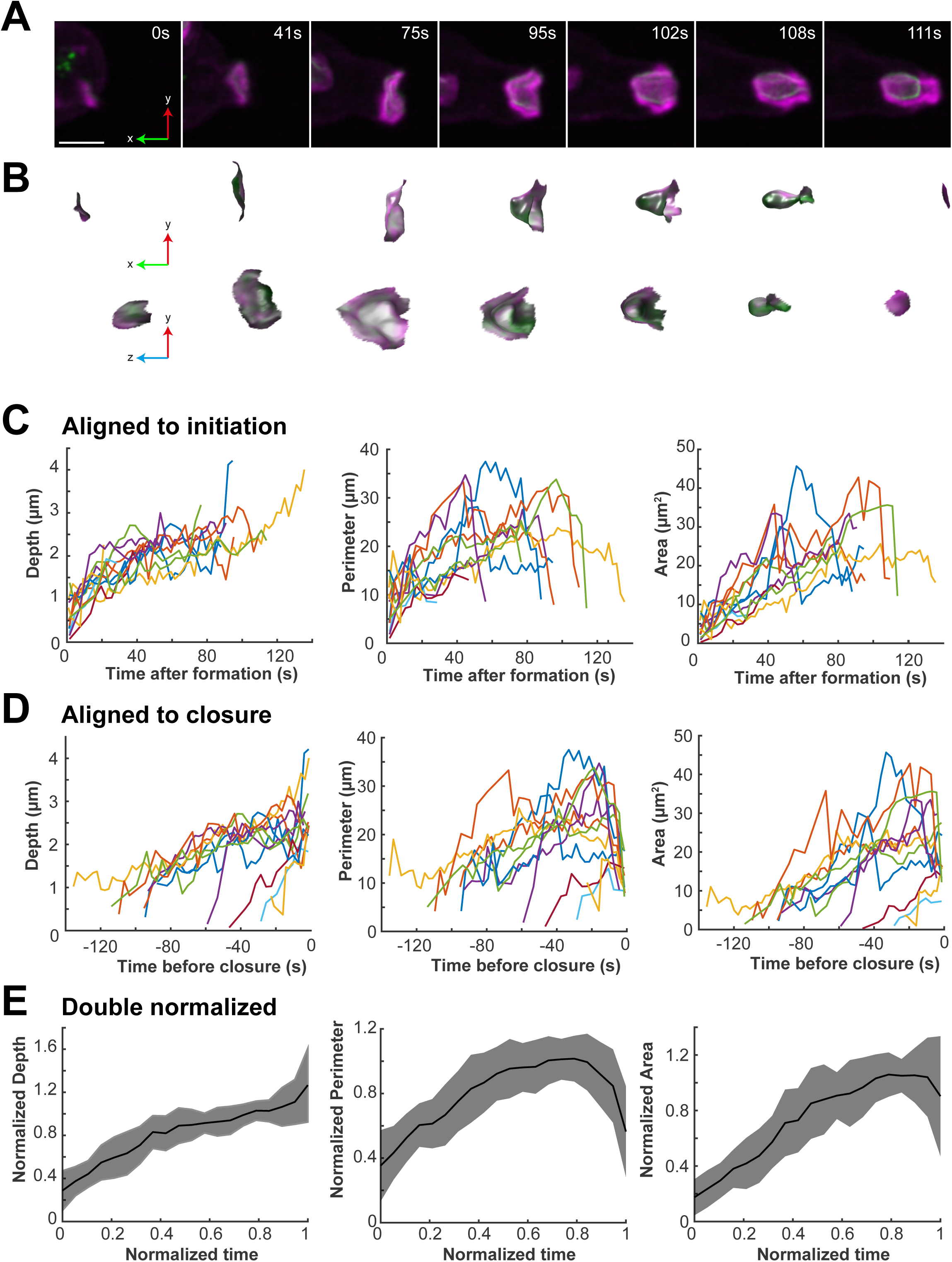
Macropinocytic cups close when their PIP3 domain stops expanding. (A) Representative domain, which originated in association with a small patch of F-actin, then expanded, deepened and closed at the lip to yield a macropinosome after 111 sec. (B) PIP3 domains corresponding to (A) dissected out computationally, shown enlarged in transverse and en face views. The macropinosome in the last frame is not visible because it is no longer attached to the cell surface. (C,D) Depth, perimeter and area of individual domains followed from origin to closure and aligned either by time of origin or of closure. Although some trends can be discerned, there is considerable variability, especially in lifetime, which varies from 30-130 sec. (E) Double normalized plots of domain depth, perimeter and area. Strikingly, domains stop expanding in area before they close. Lifetime for each domain is normalized to 1 and the dependent variable to the 75^th^ centile. The time of closure (0 sec) is taken as the last frame in which the macropinocytic cup is unclosed. PIP3 domains were selected that could be followed throughout their life, from first appearance to closure at the lip (n=11). Ax2 cells, expressing reporters for PIP3 (PkgE-PH-GFP) and F-actin (lifeAct-mCherry). Scale bars = 2 μm. See movie 14.

The life history of an average domain can be divided into phases. In the first, it expands with a steadily increasing area, while deepening and increasing in perimeter. We measured the rate of area increase in the expansion phase, both in the selected set of domains and in a larger set, obtaining values of 0.35±0.16 μm^2^ s^-1^ (N=11) and 0.30±0.34 μm^2^ s^-1^ (N=27) respectively (**Table S1**). Since the area increases approximately linearly with time, the domain boundary moves at a decreasing speed (proportional to 1/r for a circular domain). This can be regarded as a phase where an expanding wave of actin polymerization captures membrane into the cup, which remains relatively shallow and open. In the second phase, starting at around 0.6 of normalized lifetime, the domain slows and stops expanding. Finally, the perimeter deceases towards zero as the cup closes, the depth may also increase, while the area holds steady or decreases slightly. These last two phases can be regarded as a period in which a nearly constant area of membrane is manipulated by the actin cytoskeleton into a closed vesicle.

In a second test of the correlation, we selected cups which could be followed for the last minute before closure and classified them as either lip or basal closures (**Figure S4**). In lip closures the area of the domain does not increase in this period, and may even decrease just before closure, while the depth also only increases slightly, while perimeter decreases rapidly in the last 20 seconds, confirming our previous observations. Basal closure differs: although the area also does not increase over the last 30 seconds, the cup deepens around 2-fold before a macropinosome is pinched off.

These different approaches show that macropinocytic cups close when their PIP3 domain stop expanding. This suggests a mechanistic hypothesis: cup closure is caused by stalling the expansion of their PIP3 domain.

### A ‘stalled wave’ model for macropinocytic cup closure

Our observations suggest a mechanism for macropinocytic cup closure. To formalize this and test its feasibility, we built a simple model based on the following propositions **(Figure 6)**:

**Figure 6:**
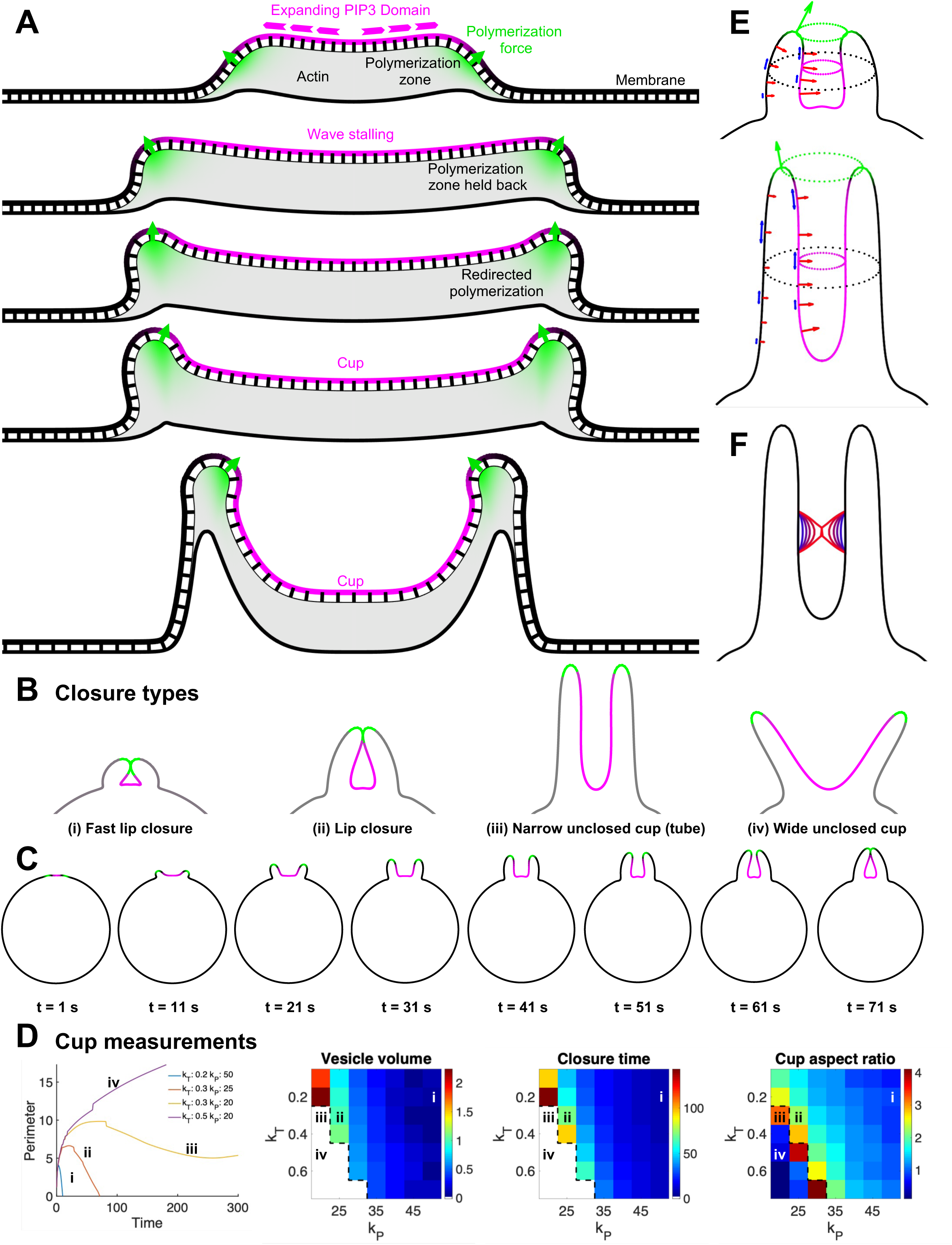
Conceptual model for macropinocytic cup expansion and closure. (A) Illustrative diagram of how the model assumptions can lead to cup formation from a PIP3 domain. Actin polymerized at the boundary is incorporated into the thick actin layer under the domain, and domain expansion pushes the polymerization zone outward to continue the process. Slowing expansion leads to actin build-up at the domain boundary, redirecting the net force upwards, allowing a cup to form. The same process guides the polymerization force inwards during lip closure, as shown in (E, top). (B) range of cup shapes generated by the model. Magenta: PIP3 domain, green: actin polymerization zone at the domain boundary. (C) Evolution of lip closure over time for the cup shown in (B, ii). (D) Cup measurements: domain perimeter for the cups in (B) over time and phase plots for volume enclosed by a surface of rotation of the domain at closure; time of closure; and aspect ratio, given by length/width of the cup at the final time point. Cups in (B) marked in all plots by numerals, k_T_: tension force coefficient, k_P_ polymerization force coefficient, black dashed lines mark the boundary between lip and no closure - only a small increase in polymerization force (25%) is required to switch from lip closure (ii) to a tube (iii). (E) Tension (red – normal, blue – tangential) and polymerization (green) forces acting on the membrane (left half only) for cups shown in (C, t = 51 s) (top) and (B, iii) (bottom), where normal tension has been modified to include curvature from the surface of rotation. The polymerization force pushes the lip inwards for the top cup and outwards for the bottom cup, due to differences in polymerization force overcoming the tension-induced contraction of the cup. Normal tension forces are similar inside both cups, (and larger than outside), but the length of the bottom cup gives more space for delamination shown in (F), giving a higher chance of base closure. (F) Delamination of the membrane from (E, bottom) moving under tension, modelled as a 3D process, which acts to minimize the surface area of the tube. Given a sufficiently large delaminated region, this process leads to closure. Lines represent the membrane at different time points, from blue (earlier) to red (later).

1. Dendritic actin polymerization occurs as a ring around the perimeter of PIP3 domains. It is protrusive and pushes against the membrane.
2. Macropinocytic cups are supported by an F-actin scaffold, to which the plasma membrane adheres. This prevents delamination and means that the membrane does not move readily slide over the scaffold and so elongation of the cup causes local increases in membrane tension
3. Cups close when expansion of the PIP3 domain stalls.

The cell is represented in the model as a 2D contour, with the rate of PIP3 expansion, force due to actin polymerization at the domain boundary, and force due to tension each controlled directly via scaling parameters, so that the consequences of different regimes can be explored. The evolving shape of macropinocytic cups is then produced by the interplay of domain expansion with forces produced by actin polymerization in the protrusive zone and membrane tension. Actin polymerization within the cup due to formins ^39^, and the signalling dynamics sustaining PIP3 domains are not considered.

**Figure 6A** shows the model intuitively: in the expansion phase (top), the PIP3 domain expands, capturing more membrane and forming an expanding F-actin wave ^40^. The zone of actin polymerization is directed outwards and does not remain under the same area of membrane for long, so that the cup remains shallow. As the domain stalls, the actin polymerization zone remains for longer under any area of membrane, the cup deepens and the zone of actin polymerization slips inside it so that it becomes directed inward, potentially giving a lip closure.

The model produces a range of behaviours depending on the parameters chosen: lip closure at different sizes; elongation with increasing membrane tension giving a basal closure if local delamination occurs; and a persistent, unclosed cup **(Figure 6B, C)**. These outcomes can be largely controlled by the polymerization force parameter (K_p_) (**Figure 6D**). The polymerization force drives the protrusion of the lip, and therefore determines the relationship between the lip and the domain boundary. Lip closure occurs if the zone of actin polymerization slips into the cup. It is prevented if the zone of actin polymerization remains near the lip, pointing outwards. This can be achieved if the tension generated in plane with the membrane (parameterized by K_t_) constricts the cup, elongating the PIP3 domain and pushing its boundary toward the lip, keeping pace with the protruding lip and maintaining the position of the polymerization zone.

In a narrow tube structure, the tension force in a 3D surface of rotation is substantially larger than in the 2D contour due to the high mean curvature of the surface (**Figure 6E**). By adapting the tension force to account for the full 3D surface, we can additionally provide a model of basal closure (**Figure 6F**). When the inward force generated by tension exceeds the adhesion force between membrane and scaffold, it causes local delamination and sealing of a macropinosome.

Our model can therefore reproduce both lip and basal closures, depending on the parameters chosen, showing that the starting propositions provide a route to understanding how macropinocytic cups form and close.

## Discussion

Lattice light-sheet microscopy gives a new window into the formation and closure of macropinocytic cups, a process that is largely unexplained. Macropinosomes in *Dictyostelium* form from cups, not from membrane flaps, and are organized around essential domains of PIP3 that extend to the lip of the cup. They are supported by a specialized F-actin framework, including actin-binding proteins, such as coronin, and the PIP3-binding myosin-1E, which is strongly concentrated within cups, and may anchor the membrane to the cytoskeleton.

Cupped or circular structures of the plasma membrane require an organizing principle to shape them. We propose that this principle is the ability of PIP3 domains to attract dendritic actin polymerization to their periphery by recruiting the Scar/WAVE and Arp2/3 complexes. Supporting this idea, VASP, which interacts with Scar/WAVE, is similarly recruited to the lips of macropinocytic cups, where it may accelerate actin polymerization ^41^. This principle may extend to other cupped structures, including large (but not small) phagosomes in *Dictyostelium* ^14, 19, 42^ and circular dorsal ruffles in mammalian cells, which similarly form around domains of PIP3 ^43^.

Unexpectedly, cups can close in two different ways: at the lip as anticipated from phagosome closure; or at the base. Basal closure has not been described before to our knowledge, though the elongated tube-like intermediates resemble the macropinocytic structures of giant amoebae ^44^. Both modes of closure appear to be driven by continued actin polymerization at the lip, which generates membrane tension within the cup: in one case we propose that the zone of actin polymerization slips inside the lip and the aperture is closed by re-directed actin polymerization; in the other, actin polymerization is not re-directed, but instead causes elongation of the cup, stretching the membrane so that eventually a macropinosome is pinched off to release the tension. Somewhat against expectation ^37, 45^ a contractile, myosin-II-based activity does not appear to be required for closure by either mechanism.

Our modelling treats macropinocytic cups in 2D and only considers mechanical forces. Yet, even within these limitations, it readily reproduces an expanding cup that closes at the lip, and one that does not close at the lip, but elongates, as in base closure. The model of Saito and Sawai ^46^, although enacted differently to our model, also incorporates the propositions of actin polymerization around PIP3 domains and domain stalling, and likewise reproduces cup formation and closure at the lip, indicating that these are robust consequences of the initial assumptions.

PIP3 domains themselves are not well understood ^47^. They can form spontaneously and are likely based on a reaction-diffusion system, with autocatalysis and cross-inhibition of diffusing molecules, including Ras, PIP3 and Rac. Active-Ras, which activates PI3-kinase, may be at the core of this system, since Ras domains can form in the absence of PIP3 domains ^18^, whereas increasing Ras activity gives bigger PIP3 domains ^19^. Their size may also be limited by the need for PI4,5P2 to diffuse into them for conversion to PIP3, as suggested by our detection of opposing gradients of these phosphoinositides. Finally, their ability to attract actin polymerization to their periphery may be due to the creation of an annulus around them, where Rac activation outweighs its inhibition ^20^.

In summary, we have described macropinocytosis in greater detail than hitherto and gained mechanistic insights, leading to a simple model for cup closure seen as resulting from stalled expansion of PIP3 domains. This provides a descriptive and conceptual framework for understanding macropinocytosis in a model organism.

## Acknowledgements

We wish to thank Dr Sharon Collier for preliminary investigations of microscopy methods, Scott Brooks for computational assistance, and Peter Devreotes and Arjan Kotholt for plasmids. We are grateful for funding from EPSRC EP/X026663/1, BBSRC BB/R004579/1 to TB; Royal Society University Research Fellowship UF140624 and URF\R\201036 and BBSRC grant BB/W006049/1 to JK; An MRC Discovery Medicine North (DiMeN) Doctoral Training Partnership (MR/N013840/1) studentship to CJM; and the Medical Research Council (part of UKRI), MRC file reference number MC_U105178783 to RRK. For purposes of open access the MRC Laboratory of Molecular Biology has applied a CC BY public copyright licence to any Author Accepted Manuscript version arising. The Warwick Lattice Light Sheet microscope and visitor programme is funded by Wellcome (208384/Z/17/Z).

## Supplementary materials

**Table S1:** Domain parameters. Domains were identified computationally as described in the Methods. Data was extracted from the stated number of movies, total elapsed time and domains. Expansion rate and geometric measurements were obtained from single domains, which did not split in the observation window. Parameters were obtained either from the set of domains used in Figure 5, which could be followed from start to finish (salmon) or larger sets, which could be followed for only the start or finish of their life (green). Domain expansion rate was calculated for the first 30 sec after *de novo* domain formation as the gradient of the change in area, calculated using a line of best fit of.

| Measurement | Mean<br>± SD | Movies<br>(n) | Time<br>total<br>(min) | Domains<br>(n) |
| --- | --- | --- | --- | --- |
| **Ax2 strain** |  |  |  |  |
| Domains per cell | 2.71±1.72 | 54 | 225 | - |
| MP formation per cell (min<sup>-1</sup>) | 1.80±1.20 | 54 | 225 | - |
| Lip closure per cell (min<sup>-1</sup>) | 1.23±0.88 | 54 | 225 | - |
| Base closure per cell (min<sup>-1</sup>) | 0.56±0.56 | 54 | 225 | - |
| Domain expansion rate* (μm<sup>2</sup> s<sup>-1</sup>) | 0.30±0.34 | 27 | 64.7 | 112 |
| Domain maximum area* (μm<sup>2</sup>) | 27.2±14.9 | 44 | 146 | 256 |
| Cup maximum depth* (μm) | 2.69±1.07 | 44 | 146 | 256 |
| Maximum domain perimeter* (μm) | 27.5±10.5 | 44 | 146 | 256 |
| Domain expansion rate** (μm<sup>2</sup> s<sup>-1</sup>) | 0.35±0.16 | 11 | 17.2 | 13 |
| Domain maximum area** (μm<sup>2</sup>) | 29.5±9.43 | 11 | 17.2 | 13 |
| Cup maximum depth** (μm) | 2.88±0.60 | 11 | 17.2 | 13 |
| Domain maximum perimeter** (μm) | 28.0±7.72 | 11 | 17.2 | 13 |
| **Other strains** |  |  |  |  |
| DdB (NF1+), MP formation per cell (min<sup>-1</sup>) | 2.52±1.47 | 2 | 12.9 | - |
| HM1200 (PI3K1-5-), MP formation per cell (min<sup>-1</sup>) | 0 | 5 | 21.1 | - |
| HM1289 (PTEN-), MP formation per cell (min<sup>-1</sup>) | 0.58±0.07 | 3 | 15.7 | - |
| RGBARG-, MP formation per cell (min<sup>-1</sup>) | 2.36±0.61 | 3 | 21.2 | - |

\*Single cups, not complex ones; \*\*Single cups used for Figure 6

**Table S2:** Reporter strains. Reporter strains were produced by transforming the indicated parental strain with the plasmid given in the first column. These plasmids are of the Paschke/Veltman series ^48, 49^, with both reporters on the same plasmid except in the case of the Nap and ArpB reporters, which were present as stable genomic insertions to which a PIP3 reporter was added.

| Plasmid | Green | Red | Parental Strain | Reporter Strain | Reference & comments |
| --- | --- | --- | --- | --- | --- |
| **Ax2 and Ax3 parental strains** |  |  |  |  |  |
| pPI304 | PIP3 | LifeAct | Ax2 | HM3478<br>HM3477 |  |
| pPI167 | Nap | PIP3 | HM1995 (Ax3) | HM3485 | REMI insertion of NAP-GFP into NAP null |
| pPI167 | ArpB | PIP3 | HM2191 (Ax2) | HM3500 | Knock-in ArpB-GFP<sup>50</sup> |
| pPI36 | Carmil | PIP3 | Ax2 | HM3498 |  |
| pPI78 | Coronin | PIP3 | Ax2 | HM3499 |  |
| pPI32 | Myo1B | PIP3 | Ax2 | HM4002 |  |
| pPI87 | Myo1E | PIP3 | Ax2 | HM4003 |  |
| pPI356 | PIP3 | PTEN | Ax2 | HM4013 |  |
| pDM1485 | MhcA | PIP3 | Ax2 | HM4021 |  |
| pJSK696 | PI3,4P2 | PIP3 | Ax2 | eJSK09 |  |
| pJSK706 | PI4,5P2 | PIP3 | Ax2 | eJSK15 |  |
| pJSK708 | PI4P | PIP3 | Ax2 | eJSK18 |  |
| pJSK709 | Tubulin | PIP3 | Ax2 | eJSK19 |  |
| **Other parental strains** |  |  |  |  |  |
| pPI304 | PIP3 | LifeAct | DdB (NF1+) | HM3481 | <sup>19</sup> |
| pPI321 | RBD | LifeAct | HM1200 (PI3K1-5-) | HM4006 | <sup>32</sup> |
| pPI304 | PIP3 | LifeAct | HM1289 (PTEN-) | HM4007 | <sup>32</sup> |
| pPI304 | PIP3 | LifeAct | RGBARG- | eJSK25 | <sup>20</sup> |

## Supplementary figures

**Figure S1: Complex, failed and non-axenic macropinocytosis**

(A, B) Closure of a complex, multi-lobed cup produces several macropinosomes almost simultaneously: (A) orthogonal views of a single Ax2 cell, the macropinocytic region of which is followed in (B). (C) Macropinocytic cup failure (arrowed) in Ax2, where after a failed attempt at closure (18.0 sec) the PIP3 domain fades away, followed by the F-actin. (D-G) Macropinocytosis is similar in non-axenic DdB cells, which have a functional NF1 RasGAP, to axenic Ax2 cells where NF1 is deleted. (D, E) closure at the lip; (F, G) closure at the base. As in Ax2 cells, DdB cells form PIP3 domains that mark macropinocytic cups, and these cups can close at lip or base. Transient tethers between the newly closed macropinosome and the cell surface also frequently form. The essential features of macropinocytosis are therefore very similar to Ax2 cells. All cells express the PIP3/lifeAct reporter combination. DdB cells were grown in HL5 supplemented with 10% FCS to stimulate growth and macropinocytosis. Scale bars B,C,D,F = 5 μm; E,G = 2 μm. See movies 4 (complex closure), 5 (failed closure), 6 (DdB, lip closure), 7 (DdB base closure).

**Figure S2: Mutants in Ras/PIP3 signalling are defective in macropinocytosis**

(A) PI3-kinase mutant (HM1200): this mutant lacks all 5 Ras-activated PI3-kinases present in the genome and has about 10% of the PIP3 and takes up less than 10% of the fluid as its parent. It still makes domains of active Ras but no macropinosomes were detected. (B, C) PTEN-mutant (HM1289) this lacks PTEN, which reverts PIP3 to PI4,5P2 and in consequence has greatly elevated PIP3 levels (∼10-fold) and in consequence elevated PIP3 across the plasm membrane. (D, E) RGBARG-mutant – this lacks a RasGAP and in consequence makes enlarged PIP3 domains, which are inefficient at macropinocytosis. Numerous small macropinosomes are formed by base closure. Representative images of each mutant expressing paired reporters for PIP3, or in the case of PI3K1-5-for active Ras (RBD - Ras Binding Domain) paired with lifeAct for F-actin. Rates of macropinosome formation are given in Table 1. Fluid uptake rates and PIP3 levels (HM1200 and HM1289) have been determined previously ^14, 18, 20, 51^. Scale bar = 5μm.

**Figure S3: Distribution of cytoskeletal reporters**

Representative images showing GFP fusion reporters (green) for the selected cytoskeletal protein with a PIP3 reporter (magenta). Coronin, carmil, myosin1B (Myo1B) and myosin1E (Myo1E) are all concentrated in macropinocytic cups, as previously described, with the PIP3-binding Myo1E closely matching the PIP3 distribution. Myosin-II (Myo-II) is not detected in macropinocytic cups at any stage but can accumulate at the contractile rear of the cell. Microtubules, marked by GFP-tubulin, move freely in the cytoplasm, and sometimes appear to touch macropinocytic cups, but no stable association is detected. Ax2 cells expressing the reporters listed in Table 2. Scale bar = 5μm.

**Figure S4: The last minute before closure of cups at lip or base**

Macropinocytic cups that could be followed for the last minute before closure were classified as either closing at the lip (19) or base (10) and the geometric parameters of the PIP3 domain computationally extracted as described in the Methods. Ax2 cells expressing the PIP3/lifeAct reporter combination. The time of closure (0 sec) is taken as the last frame in which the macropinocytic cup is unclosed.

## Movies

Movie 1: Orthogonal views of a cell producing macropinosomes by both lip and basal closure. Ax2 cell expressing reporters for F-actin and PIP3 in SUM medium. Scale bar = 5 μm.

Movie 2: segment of Movie 1 with split channels showing a macropinocytic cup closing at the lip

Movie 3: segment of Movie 1 with split channels showing a macropinocytic cup producing successive basal closures and then a final lip closure.

Movie 4: Complex macropinocytic cup giving multiple macropinosomes on closure. Ax2 cell expressing reporters for F-actin and PIP3 in SUM medium with split channels. Scale bar = 5 μm.

Movie 5: Failed macropinocytic cup. Ax2 cell expressing reporters for F-actin and PIP3 in SUM medium with split channels. Scale bar = 5 μm.

Movie 6: Macropinocytosis in strain DdB with intact NF1 – multiple lip closures. DdB cell expressing reporters for F-actin and PIP3 in SUM medium with split channels. Scale bar = 5 μm.

Movie 7: Macropinocytosis in strain DdB with intact NF1 – base followed by lip closure (left domain). DdB cell expressing reporters for F-actin and PIP3 in SUM medium with split channels. Scale bar = 5 μm.

Movie 8: Orthogonal views of a cell forming macropinosomes by both lip and basal closure. Ax3 cell expressing reporters for NAP and PIP3 in SUM medium. Scale bar = 5 μm.

Movie 9: segment of Movie 8 showing basal closure followed by lip closure. Scale bar = 2 μm.

Movie 10: Orthogonal views of a cell forming macropinosomes by both lip and basal closure. Ax2 cell expressing reporters for ArpB and PIP3 in SUM medium. Scale bar = 5 μm.

Movie 11: delamination during closure at the lip. Scale bar = 2 μm. Movie 12: delamination during closure at the base. Scale bar = 2 μm. Movie 13: delamination during a complex closure. Scale bar = 2 μm.

Movie 14: life history of PIP3 domain from origin to closure, with computationally-extracted domain shown in two orthogonal projections below. Ax2 cell expressing reporters for F-actin and PIP3 in SUM medium. Scale bar = 2 μm.

## Materials and Methods

### Cell handling

Reporter strains are listed in Table S2 and derive from Ax2(Kay) ^19^ unless otherwise indicated. They were grown at 22°C either in HL5 axenic medium (Formedium, Hunstanton, UK) in shaken suspension or on tissue-culture plates; or in association with *Klebsiella aerogenes* on SM agar plates. Non-axenic or poorly growing axenic strains, such as DdB, were grown in HL5 reinforced with 10% dialysed FCS (Gibco).

Cells were transformed with plasmids from the Paschke/Veltman series ^48, 49^, generally to express two reporters, one of which was for PIP3 (PH-PkgE ^18^). The PI45P2 reporter used was Nodulin^52^, PI34P2 was PH-TAPP1^17^ and PI4P was P4M^53^. Microtubules were imaged using GFP-a-tubulin^54^ and the F-actin reporter used was LifeAct ^55^. Cells were transformed by electroporation and selected on pre-grown bacteria with the appropriate drug ^48^ (Table S2; most plasmids are available from Addgene).

Cells for microscopy were grown for several days in HL5 medium to maximally up-regulate macropinocytosis. They were prepared for microscopy by incubation for 2-24 hr at approximately 10^5^ cells/cm^2^ in 6-well tissue culture plates in SUM low fluorescent medium (20 mM KH_2_PO_4_, 4 mM arginine, 3.7 mM glutamic acid, 8.5 mM lysine, 55 mM glucose, 2 mM MgSO_4_, 0.1 mM CaCl_2_ brought to pH 6.5 with H_3_PO_4_, plus 100 μg/ml dihydro-streptomycin) ^28^.

### Lattice light-sheet microscopy

LLSM was carried out using a 3i Lattice Light Sheet instrument with a 0.71 NA long-working-distance water-immersion excitation objective and a 1.1 NA water immersion emission objective with a total magnification of 62.5x. Images were captured on two Hamamatsu ORCA-Flash 4.0 v3 sCMOS cameras (single channel each). The sheet pattern was a Bessel lattice of 50 beams, with an inner and outer numerical aperture of 0.493 and 0.550 respectively. The thermoelectric cooler system was set to 17°C to achieve a temperature of 22°C in the sample chamber.

Before imaging, 10 μl of TetraSpeck beads (Thermofisher) were dried to a 5 mm coverslip (Mentzel Glasser #1, VWR) and mounted on the sample holder using vacuum grease (Dow Corning high vacuum grease, VWR). 10 ml of medium (SUM) was added to the bath and the sample left to equilibrate in the microscope for 30 minutes before being used for alignment and PSF measurement. Cameras were aligned by exciting the beads at both 488 nm and 560 nm and a PSF measurement taken for localization.

*Dictyostelium* cells in 6-well plates were detached by pipetting up and down and allowed to settle onto 5 mm coverslips for at least 15 minutes. A coverslip was mounted on the sample holder and transferred to the microscope containing 10 ml SUM in the bath. Samples were allowed to equilibrate before recording.

Images were recorded in two channels using 1-2% and 5-10% laser powers for 488 nm and 560 nm respectively, depending on the reporter. 3D volumes were recorded at 0.3 μm step size (0.163 μm deskewed) for 120-150 planes with a 5-10 ms exposure. This resulted in an overall speed of 2.24-3.12 seconds per volume, depending on the configuration, and an (x, y)-resolution of 0.104 μm/pixel. Volumes were recorded at this rate for 3-10 minutes.

### Processing

#### Deconvolution and registration

Because the same microscope was used for all experiments, deconvolution was made more efficient by condensing one PSF for each channel and applying it to all movies for deconvolution. Comparisons between these deconvolved images and a sample of movies deconvolved using PSFs generated immediately prior to the movies showed no substantial change in intensity but did yield a small translation of the image. This translation was accounted for by measuring cross-correlations of the channels within a range of 5 voxels (twice the largest observed translation) and selecting the translation that yielded the maximum average cross-correlation for the whole movie. Deconvolution was performed using the Richardson-Lucy method ^56^ (50 iterations).

#### Cell segmentation

All movies were segmented using the curvature-enhanced random walker ^27^. The segmentation method itself requires only two parameters. However, some pre- and postprocessing is often required to enable accurate segmentation, which increases the number of parameters.

To identify parameter configurations for the segmentation, movies assigned one of three groups: (1) those with PIP3 in the red channel, (2) those with PIP3 in the green channel without LifeAct, and (3) those with PIP3 and LifeAct. A parameter configuration that yielded accurate segmentation for most movies in each group was identified. Segmentations were manually verified, and any errors to segmentation of PIP3 patches were excluded from subsequent analysis. Parameter configurations for movies with a low segmentation accuracy of PIP3 patches were adjusted to improve the accuracy.

Movies in group (1), which contains the majority of the movies, were segmented using similar parameter configurations, with 70% using a single configuration. Similarly, group (2) mostly used the same parameter configuration (80%), with small variation from this configuration otherwise. Group (3) had the highest variation in parameter configurations, due to the fact that neither PIP3 nor actin have a strong presence in the cytoplasm leading to segmentation errors. However, there was a large overlap in parameter configurations, with at most four distinct values for any given parameter.

The segmentation error rate for each PIP3 domain was calculated by manually grading the segmented surfaces (further details below). An average of 6.97% +/-0.83% (standard error) of frames were identified as having a segmentation error and were excluded from subsequent analysis.

### Fluorescence projection

Triangulated surface meshes were obtained from the segmented movies using Matlab’s isosurface algorithm. These meshes take the form of a set of vertices and a set of triangular faces that connect the vertices. In order to measure the fluorescence on the membrane, the fluorescence values in the image were projected onto the surface. This was performed using line scans along the direction normal to the surface. For a given vertex *v* on the surface, the image voxel coordinates along a line *l* normal to the surface through *v* were calculated. Line scans were truncated to avoid including voxels containing membrane at another part of the cell surface; if a point *p* in *l* is closer to another point on the surface than to *v*, then *l* is truncated at *p* and any points beyond *p* are discarded from *l*. A minimum distance of 2 voxels (approx. 0.2 μm) was imposed on the truncation to prevent zero-length line scan in highly curved parts of the surface. Lines had a maximum length of 1 μm (1.5 μm for actin fluorescence). Each vertex was assigned the maximum fluorescence value along its line scan. Background subtraction of the fluorescence was applied to the image prior to projection using a top hat filter of radius 10 voxels for all images.

For the purpose of computing line scans, all surfaces were smoothed using Laplacian smoothing. Projecting actin fluorescence onto the surface additionally required shrinking the surface areas of high mean curvature. This is because the actin cortex lies slightly below the membrane, and therefore the peak fluorescence is farther from the surface than other markers. Given that highly curved areas introduce extreme truncation of line scans, this would lead to the actin at the lip of the macropinocytic cups being reduced. We solved this problem by shrinking the surface under mean curvature flow, which forces the surface to shrink in highly curved regions. In order to avoid simultaneously shrinking the cups during this process, mean curvature flow was only applied in regions of positive mean curvature. Positive mean curvature flow was applied until any point on the surface had moved up to a maximum of 5 μm from its original position.

### Annotation

Surfaces were annotated using MiCellAnnGELo, a VR software for rapid annotation of surfaces ^57^. Markers were placed on the surface using the marker placement tool in MiCellAnnGELo at locations within PIP3 domains in each movie. For *de novo* domain formation events, markers were placed in frames prior to this event at the approximate location of the forming domain. Similarly, after domain elimination events markers were placed in subsequent frames at the approximate location of the extinguished domain. When a single domain contained more than one cup, separate markers were placed in each cup. Markers were grouped in sequences by first measuring the distance from markers in each frame to markers in the following frame and pairing each marker to the closest marker in the following frame. This pairing process was repeated in reverse, and two markers were grouped together if they paired in both directions. Occasionally, markers moved large distances between frames, for example during a closure event, leading to them no longer pairing with the corresponding marker in the previous frame. These markers we regrouped later in the pipeline by comparing manual annotations of sequences and joining sequences with no overlapping frames, but with the same manual annotations in each frame.

### PIP3 domain segmentation

Domains were identified around each marker using Otsu thresholding ^58^ of the PIP3 fluorescence values on the surface. Threshold values were computed for each marker using the histogram from vertices nearest to the given marker in the geodesic sense. In cases where structures such as cup lips or ruffles present as thin protrusions with PIP3 on only one side of the structure, a low level of PIP3 will be projected onto the other side in the surface representation due to spreading of the fluorescence during image capture. In order to avoid including the opposite side of such structures in the domain, mean curvature is used to separate the two sides, while also providing geometric definitions for *cups* and *lips*.

For a given marker, a *cup* is defined as a connected area on the surface about the marker, where all vertices have PIP3 intensity above the Otsu threshold and mean curvature less than 0 (concave). The lip is defined as the band of vertices, grouped by geodesic distance from the cup boundary, with the highest average mean curvature (highly convex). The *domain* about the given marker is the set of all vertices within the lip contour with PIP3 value above the Otsu threshold. In cases where *cups, domains*, or *lips* for two markers intersect, it is necessary to further segment these structures to maintain correspondence with the markers. This segmentation was performed by dividing the structures along the line of equal geodesic distance between the two markers. *Domains* that required segmentation in this manner were automatically graded as being split *domains*. For cases where the PIP3 intensity was below the threshold value at the marker, the marker vertex and all vertices sharing a face with that vertex were marked as the *domain* for ease of processing. The frame when this false *domain* changes to a true *domain* were automatically marked as *domain* formation events, and the reverse as *domain* elimination events.

Each of the *cups, domains*, and *lips* were additionally smoothed by dilating (adding neighbouring vertices, applied twice), filling (adding connected components surrounded by the vertices of the given feature in the mesh graph), and eroding (the reverse of the dilation operation).

### PIP3 domain grading

Each domain was subsequently graded manually for each frame to mark the events of *De novo* domain formation, domain elimination, lip closure, base closure, failed closure, and domain splitting. Domains were also assessed for cell segmentation and domain segmentation errors, which were noted and excluded from subsequent analysis. The automated grading above provided an initial set of gradings, which were adjusted where necessary.

Gradings were made by visual comparison of the surface and the 3D maximum intensity projection of the movie. A custom Jupyter notebook was designed to allow rapid grading of domains, using ITKWidgets with PyVista to render the surfaces ^59^. This notebook allows the user to change surface colours between fluorescence and domains, as well as providing controls for opacity and colour scaling. Buttons for grading allow the marking of base, lip, and failed closures, as well as domain splitting, segmentation and domain detection errors, and noting the start and end of the domain.

For a given marker, frames between domain start and end gradings are taken to be the times when the domain is present. For domains with no start frame (either formed prior to the start of the movie or formed from splitting), the domain is assumed to start from the first frame that the corresponding marker is present. Similarly, domains with no end frame are assumed to end when the marker is no longer present. Frames where the domain was not automatically detected for a given marker (see previous section) but manually marked as having a domain (and vice versa) were automatically graded as having a domain segmentation error.

In the subsequent analysis, measurements of a domain graded as split or to have an error in a given frame were excluded for that frame. Where time-dependent measurements were taken, domains which were graded as split at any time in the interval being measured were excluded.

### Measurements

#### Fluorescence measurements

Fluorescence measurements were made relative to the domain boundary. Additionally, the mean curvature of the surface was computed and processed in this manner, without the background subtraction step. The surface outside and inside this boundary was partitioned by geodesic distance to the boundary, where bands outside the domain were equally spaced at 0.2 μm, and bands inside the domains were equally spaced to 0.1*domain depth (defined below). Using a normalized distance inside the domain allows comparison between domains of different sizes. The average fluorescence values were taken in each band, giving a 1-dimensional profile of the fluorescence values for each time point. For cells with multiple domains, bands around a domain were restricted to points on the surface closest to the given domain’s boundary in the geodesic sense.

For each domain, the background fluorescence measurement in each frame was taken to be the average fluorescence 5 μm outside the domain. The median background fluorescence over the lifetime of the domain was taken to be the overall background. Fluorescence values were normalized by dividing by this background value (fold normalization).

To give an overall profile of fluorescence and curvature, the average normalized 1-dimensional profile was taken for each domain, and these averages aggregated to give the plots shown in Fig 2.

For time-dependent analysis, closure events were taken to be the temporal reference points. The number of domain sequences was reduced to include only those that contained the closure event being investigated (either lip or base closure), and the required time interval preceding the event. Due to variable time steps of the movies and some frames being rejected in the quality control step above, fluorescence values were linearly interpolated at every 2.5 seconds prior to the closure time. The average interpolated 1-dimensional fluorescence profiles for each time point were thus computed and plotted as colour values on a 2D image. The image was distorted to account for distance normalization of the profiles by scaling the part of the image representing points inside the domain by the average domain depth along the distance axis.

#### PIP3 domain geometry

Geometric measurements for each domain were made using the surface geometry: the perimeter was taken to be the total length of edges along the boundary of the domain; the area was given by the total area of all faces in the domain, and the depth was taken to be the maximum distance of any point in the domain to the domain boundary.

In order to compare the geometry of multiple domains, two approaches were used. In the first, all domains captured undergoing *de novo* formation, lip closure, and elimination were aggregated by normalizing in both time and the geometric measurement, allowing the observation of general trends in the geometry (Figure 5). Time normalization was achieved by setting the formation time to 0 and closure time to 1 and interpolating values linearly at 20 evenly spaced time points between these two events. Measurements were normalized by dividing by the 75^th^ percentile. In the second comparison, measurements for the last minute prior to closure (Figure S4) were aggregated using the closure time as a reference point. Due to the variable time between frames and some frames being excluded during quality control, values were linearly interpolated to obtain a series of values uniformly spaced in time for each dataset, which could then be aggregated at each time point. An interval time of 2.5 s was taken, since this is close to the frame interval in most of the movies. The reference time for each event was calculated as halfway between the frame containing the event and the previous frame.

### Visualization

#### 3D maximum intensity projections

It is necessary in the above to visually compare 3D surface data with 3D volume data. To enable rapid visual inspection of cells, 3-way maximum intensity projections were generated for each frame by taking the projections in each of x, y, and z after rescaling in z to produce isotropic resolution. These images were combined into one as a montage, placing the projection in z (represented in the (x, y)-plane) in the top left, the projection in y ((x, z)-plane) in the bottom left, and the projection in x ((y, z)-plane) in the top right. This yielded a view where the projections with lower resolution (x and y) could be easily compared to the high-resolution projection (z) due to the alignment of the common axes. These projections allowed easier identification of closure events than the projection in z alone because they provide some of the 3D information lacking in the single projection.

#### Surfaces

Surface rendering for purely visualization purposes was performed in Matlab (see above for annotation using VR and grading using ITKWidgets in Jupyter). The surfaces were preprocessed by applying Laplacian smoothing (3 iterations), to provide a smoother surface to view. A customized Matlab live script was used to allow fast rendering of the surfaces. Surface colour and opacity can be linked independently to fluorescence, curvature, or PIP3 patches. These values can be scaled to allow better visualization of the surface for the length of the movie.

#### Modelling

A mathematical model is used to simulate the behaviour of cells during macropinocytosis. Our aim with this model is to show the feasibility of cup formation and closure resulting from a polymerization force acting at the boundary of a PIP3 domain, given that the domain area expansion stalls at some time point.

The model is based on a previous contour model used to study blebbing ^60^. The total force, F_T_, at a given vertex is

F_T_ = F_tension_ + F_bending_ + F_area_ + F_poly,_

where F_tension_ is the force due to membrane tension, modelled as elastic stress induced by stretching the contour, F_bending_ is the force due to bending of the membrane, F_area_ is a force preserving the area inside the contour, and F_poly_ is the normal force due to actin polymerization. The first two of these forces are calculated in the same manner as previously ^60^, F_area_=k_A_(A/A_0_-1) for area inside the contour A with initial value A_0_ and force coefficient k_A_, and F_poly_ is applied along the normal direction at the domain boundary as described below. Bending and area force coefficients are kept constant in all simulations.

While a contour model is 2D in nature, we may interpret it as approximating a slice through a cell with rotational symmetry, allowing us to calculate values for the perimeter (Figure 6D) from 2D measurements.

We define a PIP3 domain on the contour as a portion of the contour with length d, expanding at a rate v_P_/d for a constant value v_P_. This gives a slowing rate of expansion in length, while the domain area, calculated using a surface of rotation, stops expanding when approaching lip closure due to the shape changes of the domain.

The contour is initiated as a circle with radius 5 μm, discretized using 1000 line segments, and PIP3 patch of initial width 0.02 μm. Throughout the simulation, the force F_poly_ is applied as a constant multiple, k_P_, of a Gaussian function centered at the domain boundary on both sides with standard deviation 0.25 μm. A domain expansion speed of 0.1 / d μm s^-1^ for domain width d is used for all experiments.

#### Base closure

This 2D model can produce range of behaviours of cups (Figure 6B). If we also consider the 3D membrane tension force within a cup (Figure 6E) and assume a rigid actin scaffold is holding the membrane in place, we can identify a potential mechanism for base closure, given a partial delamination of the membrane from the actin scaffold (Figure 6F).

If a portion of the membrane in the cup, approximating a cylinder of length l and width w, delaminates from the actin cytoskeleton, then it will start to move under membrane tension. Here, we model the tension force as proportional to mean curvature, while assuming all other forces are negligible. If the aspect ratio l/w < 0.66 (approx.), then the membrane in the cylinder can attain a minimum surface area without closure. We therefore enforce an aspect ratio greater than 0.66 ^61^ when setting the length of the delamination zone, allowing the membrane to continue to move inwards until closure. It is conceivable that smaller initial delamination lengths could also lead to base closure, since tension at the edges of the delaminated region would be increased, leading to further detachment from the actin cytoskeleton. A similar effect can also occur during lip closure but is not essential for closure in this case.

## References

1. Swanson, J.A. Shaping cups into phagosomes and macropinosomes. Nature Rev Molec Cell Biol 9, 639–649 (2008).

2. King, J.S. & Kay, R.R. The origins and evolution of macropinocytosis. Philos Trans R Soc Lond B Biol Sci 374, 20180158 (2019).

3. Stow, J.L., Hung, Y. & Wall, A.A. Macropinocytosis: Insights from immunology and cancer. Current opinion in cell biology 65, 131–140 (2020).

4. Mylvaganam, S., Freeman, S.A. & Grinstein, S. The cytoskeleton in phagocytosis and macropinocytosis. Current Biology 31, R619–R632 (2021).

5. Commisso, C. et al. Macropinocytosis of protein is an amino acid supply route in Ras-transformed cells. Nature 497, 633–637 (2013).

6. Hacker, U., Albrecht, R. & Maniak, M. Fluid-phase uptake by macropinocytosis in Dictyostelium. J Cell Sci 110, 105–112 (1997).

7. Sallusto, F., Cella, M., Danieli, C. & Lanzavecchia, A. Dendritic Cells Use Macropinocytosis and the Mannose Receptor to Concentrate Macromolecules in the Major Histocompatibility Complex Class-Ii Compartment - down-Regulation by Cytokines and Bacterial Products. Journal of Experimental Medicine 182, 389–400 (1995).

8. Mercer, J. & Helenius, A. Gulping rather than sipping: macropinocytosis as a way of virus entry. Current opinion in microbiology 15, 490–499 (2012).

9. Kranz, L.M. et al. Systemic RNA delivery to dendritic cells exploits antiviral defence for cancer immunotherapy. Nature 534, 396-+ (2016).

10. Lewis, W.H. Pinocytosis. Bull Johns Hopkins Hosp 49, 17–27 (1931).

11. Swanson, J.A. & King, J.S. The breadth of macropinocytosis research. Philos Trans R Soc Lond B Biol Sci 374, 20180146 (2019).

12. Araki, N., Johnson, M.T. & Swanson, J.A. A role for phosphoinositide 3-kinase in the completion of macropinocytosis and phagocytosis by macrophages. J Cell Biol 135, 1249–1260 (1996).

13. Buczynski, G. et al. Inactivation of two Dictyostelium discoideum genes, DdPIK1 and DdPIK2, encoding proteins related to mammalian phosphatidylinositide 3-kinases, results in defects in endocytosis, lysosome to postlysosome transport, and actin cytoskeleton organization. J Cell Biol 136, 1271–1286 (1997).

14. Hoeller, O. et al. Two distinct functions for PI3-kinases in macropinocytosis. J Cell Sci 126, 4296–4307 (2013).

15. Yoshida, S., Hoppe, A.D., Araki, N. & Swanson, J.A. Sequential signaling in plasma-membrane domains during macropinosome formation in macrophages. J Cell Sci 122, 3250–3261 (2009).

16. Parent, C.A., Blacklock, B.J., Froelich, W.M., Murphy, D.B. & Devreotes, P.N. G Protein signaling events are activated at the leading edge of chemotactic cells. Cell 95, 81–91 (1998).

17. Dormann, D., Weijer, G., Dowler, S. & Weijer, C.J. In vivo analysis of 3-phosphoinositide dynamics during Dictyostelium phagocytosis and chemotaxis. J Cell Sci 117, 6497–6509 (2004).

18. Veltman, D.M. et al. A plasma membrane template for macropinocytic cups. eLife 5: e20085 doi: 10.7554/eLife.20085 (2016).

19. Bloomfield, G. et al. Neurofibromin controls macropinocytosis and phagocytosis in Dictyostelium. eLife 4: e04940 doi: 10.7554/eLife.04940 (2015).

20. Buckley, C.M. et al. Coordinated Ras and Rac Activity Shapes Macropinocytic Cups and Enables Phagocytosis of Geometrically Diverse Bacteria. Current Biology 30, 2912–2926 (2020).

21. Marinovic, M. et al. IQGAP-related protein IqgC suppresses Ras signaling during large-scale endocytosis. Proc Natl Acad Sci U S A 116, 1289–1298 (2019).

22. Williams, T.D., Peak-Chew, S.Y., Paschke, P. & Kay, R.R. Akt and SGK protein kinases are required for efficient feeding by macropinocytosis. J Cell Sci 132, jcs.224998 (2019).

23. Yang, Y.H. et al. Leep1 interacts with PIP3 and the Scar/WAVE complex to regulate cell migration and macropinocytosis. Journal of Cell Biology 220 (2021).

24. Chen, B.C. et al. Lattice light-sheet microscopy: imaging molecules to embryos at high spatiotemporal resolution. Science 346, 1257998 (2014).

25. Condon, N.D. et al. Macropinosome formation by tent pole ruffling in macrophages. J Cell Biol 217, 3873–3885 (2018).

26. Quinn, S.E. et al. The structural dynamics of macropinosome formation and PI3-kinase-mediated sealing revealed by lattice light sheet microscopy. Nature communications 12 (2021).

27. Lutton, J.E., Collier, S. & Bretschneider, T. A Curvature-Enhanced Random Walker Segmentation Method for Detailed Capture of 3D Cell Surface Membranes. Ieee T Med Imaging 40, 514–526 (2021).

28. Williams, T.D. & Kay, R.R. The physiological regulation of macropinocytosis during Dictyostelium growth and development. J Cell Sci 131, jcs.213736 (2018).

29. de Hostos, E.L., Bradtke, B., Lottspeich, F., Guggenheim, R. & Gerisch, G. Coronin, an actin binding protein of Dictyostelium discoideum localized to cell surface projections, has sequence similarities to G-protein beta-subunits. Embo J 10, 4097–4104 (1991).

30. Bloomfield, G., Tanaka, Y., Skelton, J., Ivens, A. & Kay, R.R. Widespread duplications in the genomes of laboratory stocks of Dictyostelium discoideum. Genome Biol. 9, R75 (2008).

31. Iijima, M. & Devreotes, P. Tumor suppressor PTEN mediates sensing of chemoattractant gradients. Cell 109, 599–610 (2002).

32. Hoeller, O. & Kay, R.R. Chemotaxis in the absence of PIP3 gradients. Curr. Biol. 17, 813–817 (2007).

33. Chan, K.T., Creed, S.J. & Bear, J.E. Unraveling the enigma: progress towards understanding the coronin family of actin regulators. Trends Cell Biol 21, 481–488 (2011).

34. McConnell, R.E. & Tyska, M.J. Leveraging the membrane - cytoskeleton interface with myosin-1. Trends Cell Biol 20, 418–426 (2010).

35. Brzeska, H., Koech, H., Pridham, K.J., Korn, E.D. & Titus, M.A. Selective localization of myosin-I proteins in macropinosomes and actin waves. Cytoskeleton (Hoboken) 73, 68–82 (2016).

36. Jung, G., Remmert, K., Wu, X.F., Volosky, J.M. & Hammer, J.A. The Dictyostelium CARMIL protein links capping protein and the Arp2/3 complex to type I myosins through their SH3 domains. J Cell Biol 153, 1479–1497 (2001).

37. Swanson, J.A. et al. A contractile activity that closes phagosomes in macrophages. J Cell Sci 112 (Pt 3), 307–316 (1999).

38. Shu, S., Liu, X. & Korn, E.D. Blebbistatin and blebbistatin-inactivated myosin II inhibit myosin II-independent processes in Dictyostelium. Proc Natl Acad Sci U S A 102, 1472–1477 (2005).

39. Junemann, A. et al. A Diaphanous-related formin links Ras signaling directly to actin assembly in macropinocytosis and phagocytosis. Proc Natl Acad Sci U S A 113, 7464–7473 (2016).

40. Bretschneider, T. et al. The three-dimensional dynamics of actin waves, a model of cytoskeletal self-organization. Biophys J. 96, 2888–2900 (2009).

41. Korber, S. & Faix, J. VASP boosts protrusive activity of macroendocytic cups and drives phagosome rocketing after internalization. Eur J Cell Biol 101 (2022).

42. Clarke, M. et al. Curvature recognition and force generation in phagocytosis. Bmc Biol 8, 154 (2010).

43. Yoshida, S., Pacitto, R., Sesi, C., Kotula, L. & Swanson, J.A. Dorsal ruffles enhance activation of Akt by growth factors. J Cell Sci 131, jcs.220517 (2018).

44. Chapman-Andresen, C. Endocytosis in freshwater amebas. Physiological reviews 57, 371–385 (1977).

45. Araki, N., Hatae, T., Furukawa, A. & Swanson, J.A. Phosphoinositide-3-kinase-independent contractile activities associated with Fc gamma-receptor-mediated phagocytosis and macropinocytosis in macrophages. J Cell Sci 116, 247–257 (2003).

46. Saito, N. & Sawai, S. Three-dimensional morphodynamic simulations of macropinocytic cups. Iscience 24 (2021).

47. Kay, R.R., Williams, T.D., Manton, J.D., Traynor, D. & Paschke, P. Living on soup: macropinocytic feeding in amoebae. International Journal of Developmental Biology 63, 473–483 (2019).

48. Paschke, P. et al. Rapid and efficient genetic engineering of both wild type and axenic strains of Dictyostelium discoideum. PLoS One 13, e0196809 (2018).

49. Veltman, D.M., Akar, G., Bosgraaf, L. & Van Haastert, P.J. A new set of small, extrachromosomal expression vectors for Dictyostelium discoideum. Plasmid 61, 110–118 (2009).

50. Langridge, P.D. & Kay, R.R. Mutants in the Dictyostelium Arp2/3 complex and chemoattractant-induced actin polymerization. Exptl Cell Res 313, 2563–2574 (2007).

51. Clark, J. et al. Dictyostelium uses ether-linked inositol phospholipids for intracellular signalling. Embo J 33, 2188–2200 (2014).

52. Miao, Y. et al. Altering the threshold of an excitable signal transduction network changes cell migratory modes. Nat Cell Biol 19, 329–340 (2017).

53. Brombacher, E. et al. Rab1 guanine nucleotide exchange factor SidM is a major phosphatidylinositol 4-phosphate-binding effector protein of Legionella pneumophila. J Biol Chem 284, 4846–4856 (2009).

54. King, J.S., Veltman, D.M., Georgiou, M., Baum, B. & Insall, R.H. SCAR/WAVE is activated at mitosis and drives myosin-independent cytokinesis. J Cell Sci 123, 2246–2255 (2010).

55. Riedl, J. et al. Lifeact: a versatile marker to visualize F-actin. Nature methods 5, 605–607 (2008).

56. Richardson, W.H. Bayesian-Based Iterative Method of Image Restoration. J Opt Soc Am 62, 55-+ (1972).

57. Platt, A., Lutton, J.E. & Bretschneider, T. MiCellAnnGELo: Annotate microscopy time series of complex cell surfaces with 3D Virtual Reality. arXiv 2209.11672v1, 1–2 (2022).

58. Otsu, N. Threshold Selection Method from Gray-Level Histograms. Ieee T Syst Man Cyb 9, 62–66 (1979).

59. Sullivan, C. & Kaszynski, A. PyVista: 3D plotting and mesh analysis through a streamlined interface for the Visualization Toolkit (VTK). Journal of Open Source Software 4, 1450 (2019).

60. Collier, S., Paschke, P., Kay, R.R. & Bretschneider, T. Image based modeling of bleb site selection. Scientific reports 7, 6692 (2017).

61. Cryer, S.A. & Steen, P.H. Collapse of the Soap-Film Bridge - Quasi-Static Description. J Colloid Interf Sci 154, 276–288 (1992).

