## Supplementary figures and images for "The formation and closure of macropinocytic cups in a model system"

### Supplementary Figure S1

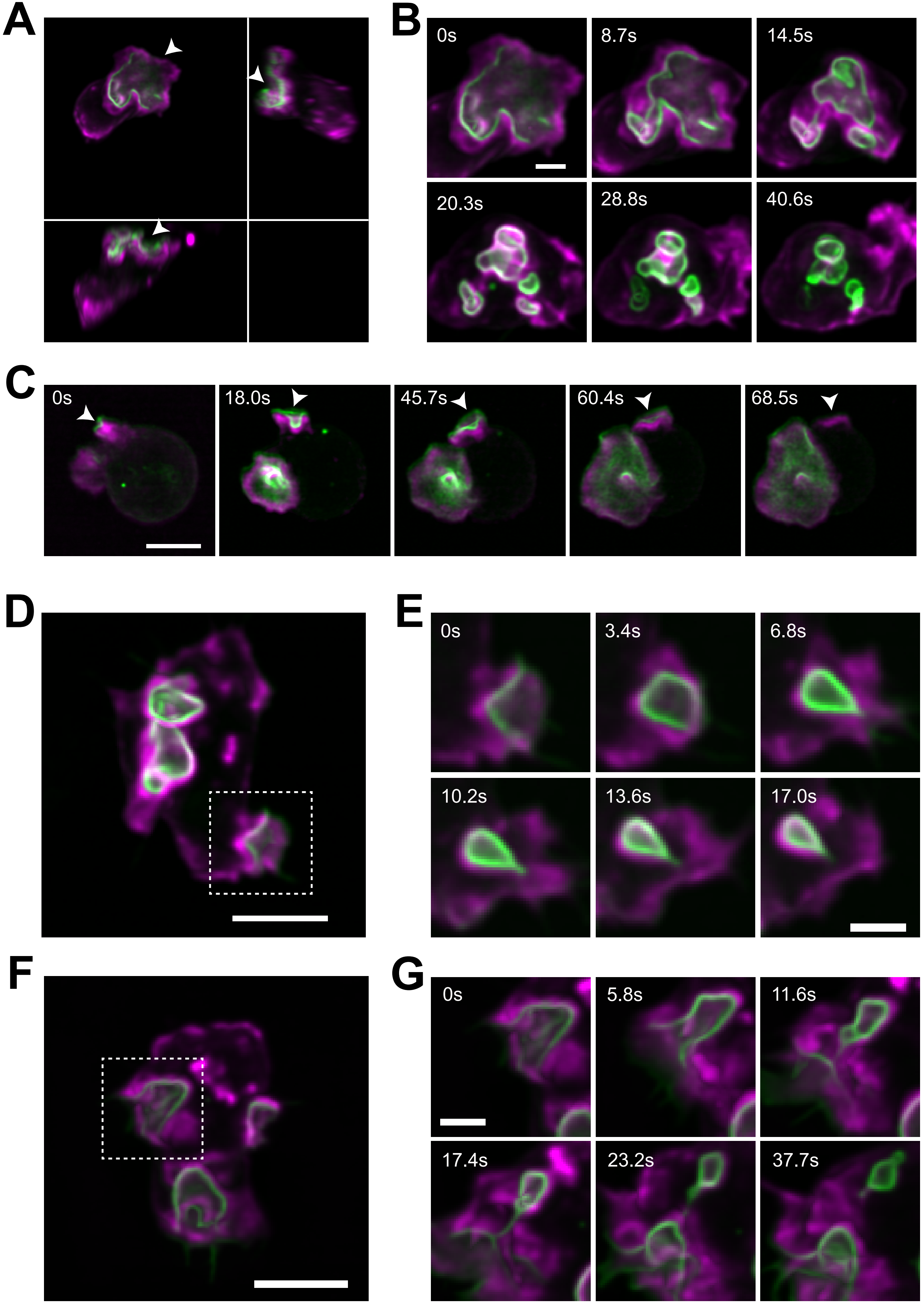

### Supplementary Figure S2

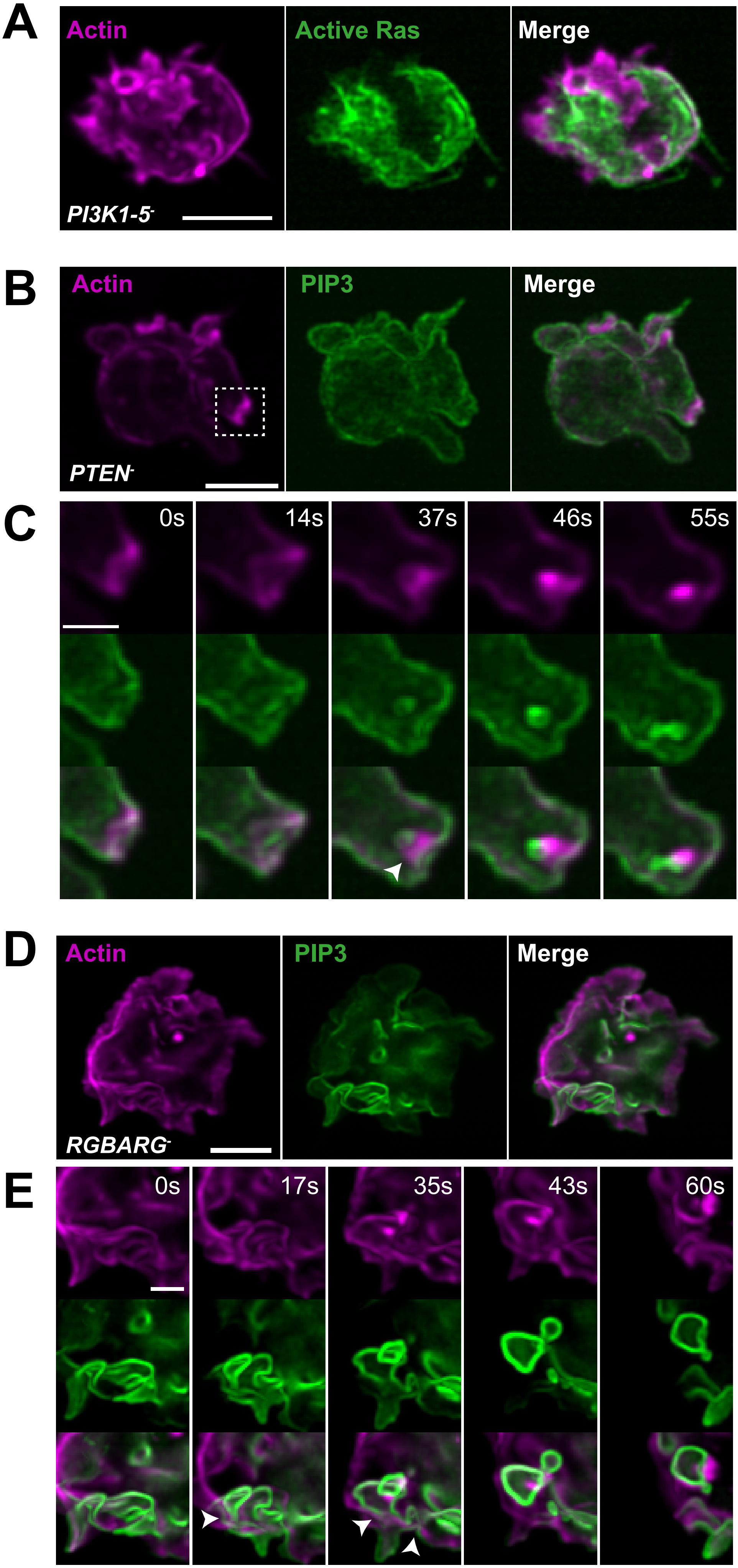

### Supplementary Figure S3

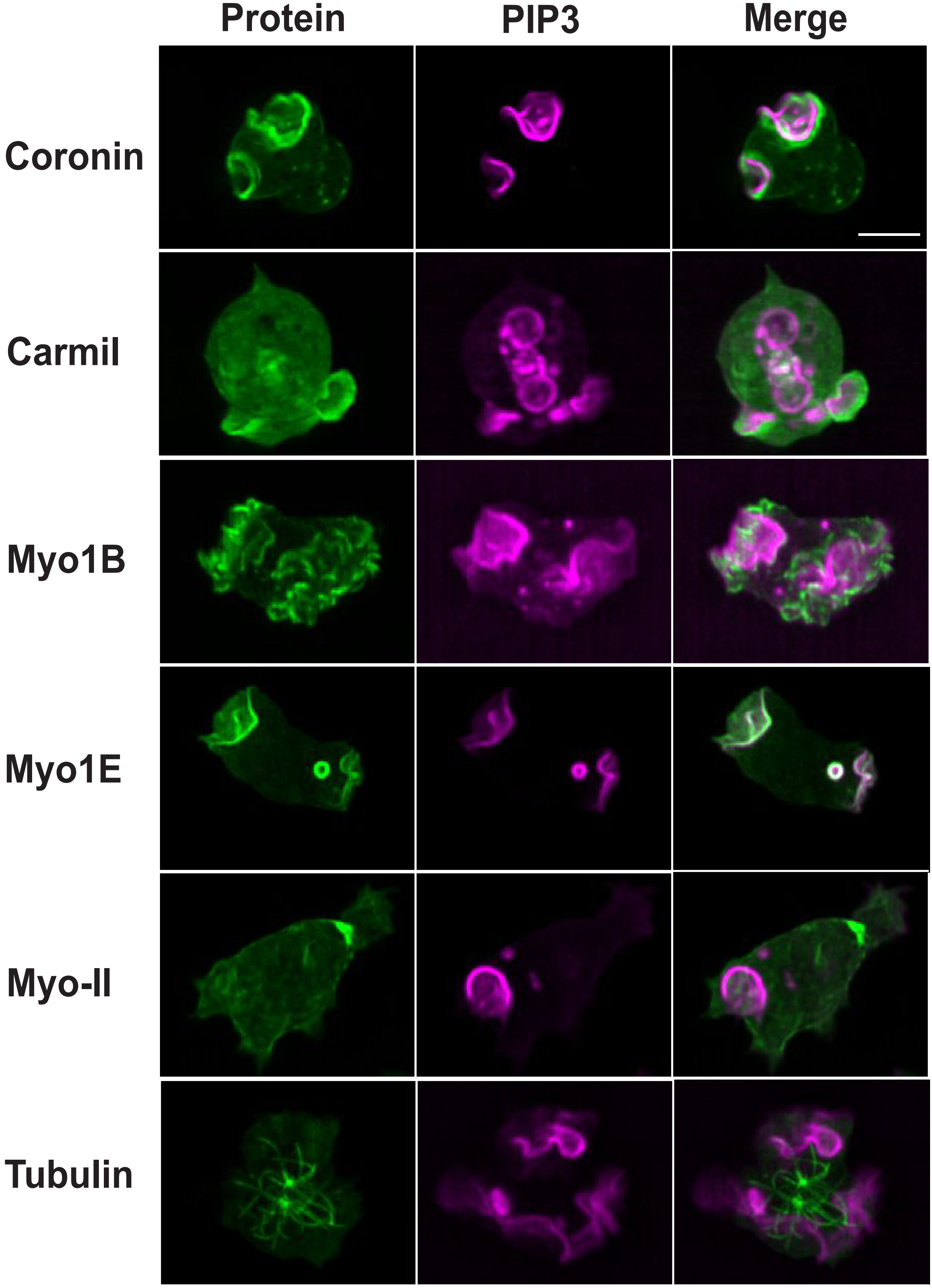

### Supplementary Figure S4

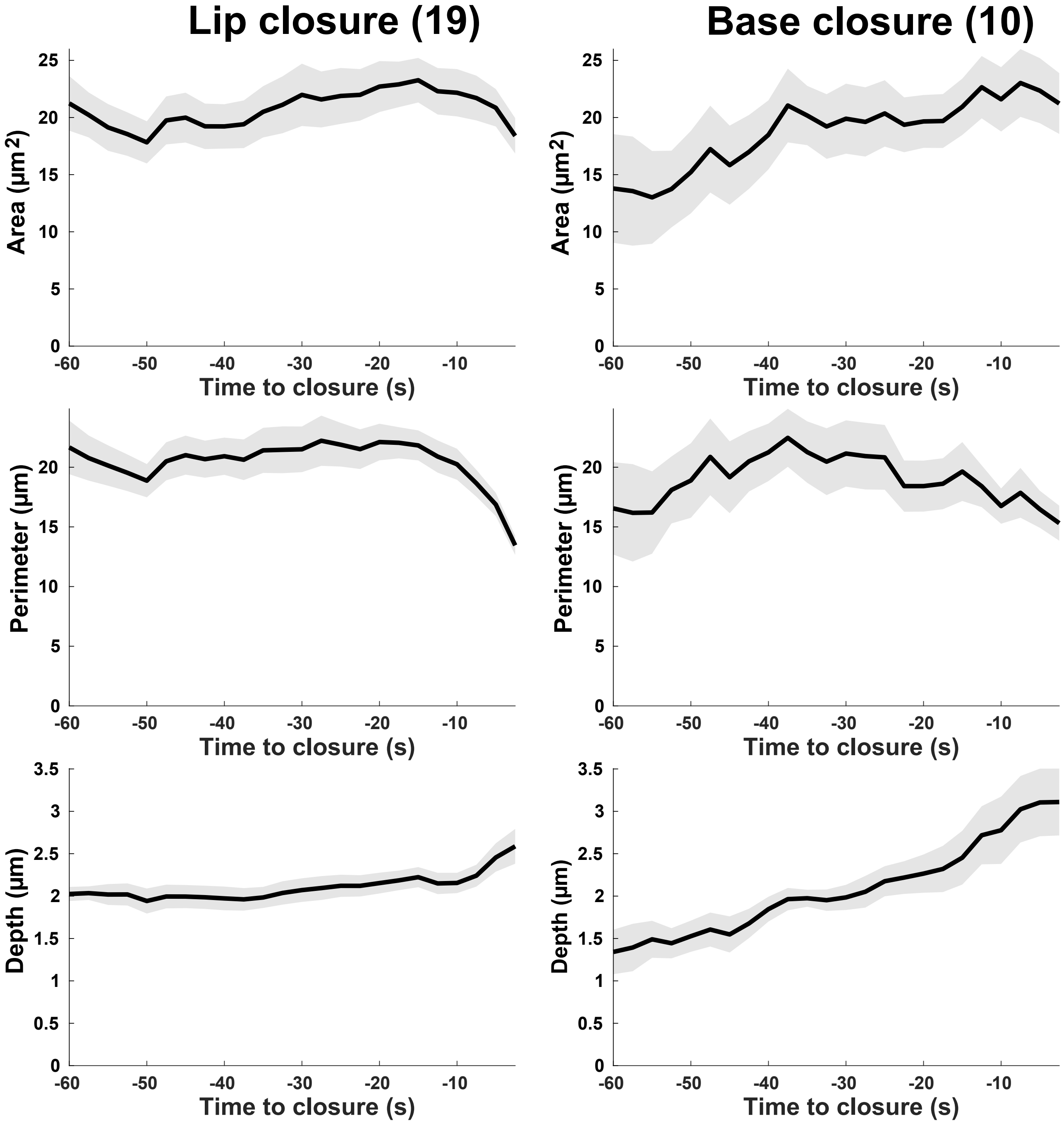
